# Acetaminophen Overdose Reveals Protease-Activated Receptor 4 as a Low-Expressing but Potent Receptor on the Hepatic Endothelium

**DOI:** 10.1101/2024.06.07.598028

**Authors:** Rahul Rajala, Audrey C.A. Cleuren, Courtney T. Griffin

## Abstract

**Background & Aims:** Hepatic endothelial cell (EC) dysfunction and centrilobular hepatocyte necrosis occur with acetaminophen (APAP) overdose. The protease thrombin, which is acutely generated during APAP overdose, can signal through protease-activated receptors 1 and 4 (PAR1/PAR4). PAR1 is a high-affinity thrombin receptor that is known to signal on ECs, whereas PAR4 is a low-affinity thrombin receptor, and evidence for its expression and function on ECs is mixed. This study aims to exploit the high levels of thrombin generated during APAP overdose to determine (1) if hepatic endothelial PAR4 is a functional receptor, and (2) endothelial-specific functions for PAR1 and PAR4 in a high thrombin setting.

**Methods:** We generated mice with conditional deletion(s) of *Par1/Par4* in ECs and overdosed them with APAP. Hepatic vascular permeability, erythrocyte congestion/bleeding, and liver function were assessed following overdose. Additionally, we investigated the expression levels of endothelial PARs and how they influence transcription in APAP-overdosed liver ECs using endothelial Translating Ribosome Affinity Purification followed by next-generation sequencing (TRAPseq).

**Results:** We found that mice deficient in high-expressing endothelial *Par1* or low-expressing *Par4* had equivalent reductions in APAP-induced hepatic vascular instability but no effect on hepatocyte necrosis. Additionally, mice with loss of endothelial *Par1* and *Par4* had reduced permeability at an earlier time point after APAP overdose when compared to mice singly deficient in either receptor in ECs. We also found that endothelial PAR1—but not PAR4—can regulate transcription in hepatic ECs.

**Conclusions:** Low-expressing PAR4 can react similarly to high-expressing PAR1 in APAP-overdosed hepatic ECs, demonstrating that PAR4 is a potent thrombin receptor. Additionally, these receptors are functionally redundant but act divergently in their expression and ability to influence transcription in hepatic ECs.

**NOMENCLATURE:** *F2r* and *F2rl3* are the gene names for PAR1 and PAR4, respectively. For simplicity, we hereafter refer to these genes as *Par1* and *Par4*.

## INTRODUCTION

The vascular endothelium, which forms the inner lining of blood vessels, is a sprawling cell layer covering a total surface area of 270-720 square meters^1^. Endothelial cells (ECs) participate in the regulation of tissue function and structure, the trafficking of hormones and nutrients^2^, the control of inflammation^3^, and the regulation of vascular permeability ^4^. With these critical and diverse roles, it is not surprising that endothelial dysfunction amplifies various pathological insults in specific organs^4–6^.

Protease-activated receptors (PARs) are a family of G protein-coupled receptors (GPCRs) with four members (PAR1-4). PAR1 and PAR4 are thrombin receptors that have been linked to many physiological processes and that are associated with pathologies like diabetes, Alzheimer’s disease, and cancer when dysregulated^7^. These receptors have different affinities for thrombin (PAR1: 50 pM and PAR4: 5000 pM)^8^, as well as distinct mechanisms for signaling^9–11^ and desensitization^7,12^. PARs can act synergistically or redundantly and can directly dimerize to enact a biased signaling response^13^. Both PAR1 and PAR4 are expressed on ECs; however, endothelial PAR4 is expressed at low levels, so there is a paucity of data on its function^14^. For example, since the receptor was reported to be present and functional on mouse ECs in the early 2000s^15,16^, only about 90 subsequent papers about endothelial PAR4 have been published. Among those papers, approximately one in six has reported that PAR4 is non-functional on ECs. Some of this controversy about the functionality of endothelial PAR4 may be explained by its low or undetectable expression on cultured ECs, including primary human umbilical vein ECs (HUVECs) that are broadly used for EC research. The lack of suitable *in vitro* tools to study endothelial PAR4 has led to the prevailing theory that PAR4 is an inconsequential thrombin receptor on ECs. However, knowing that cultured ECs can undergo phenotypic drift that results in divergent gene expression patterns from *in vivo* ECs^17,18^, we recognized a need to assess endothelial PAR4 function *in vivo* with cell-specific gene deletions in a setting of high thrombin generation.

In this study, we chose acetaminophen (APAP) overdose as a model for exploring endothelial PAR4 and PAR1 functionality. APAP overdose is the leading cause of acute liver injury in the United States^19^. The pathology of APAP is characterized by centrilobular hepatocyte necrosis and congestion of sinusoidal blood vessels^19^. Importantly for this study, thrombin levels are significantly increased during APAP overdose, making this an ideal model to study the function of endothelial thrombin receptors, and both PAR1 and PAR4 have been shown to contribute to APAP pathology in mice^20,21^. However, previous studies have used global knockout (KO) mice to study PAR1 and PAR4 in the context of APAP overdose, which makes it difficult to dissect cell-specific roles for these receptors in the hepatic niche. For example, *Par4^-/-^* mice have less liver damage than wildtype mice at 6 and 24 hours after APAP overdose, as indicated by reduced hepatocyte necrosis and plasma alanine aminotransferase (ALT) activity^21^. However, when bone marrow from *Par4^-/-^* deficient mice was transplanted into irradiated wildtype mice, the beneficial reduction in hepatocyte necrosis and ALT activity was lost^21^. The authors of this study speculated that the cell-specific loss of PAR4 on ECs—rather than on platelets, where it is robustly expressed—was responsible for the reduced hepatocyte necrosis seen in *Par4^-/-^* mice after APAP overdose^15^. Similarly, *Par1^-/-^*mice have reduced plasma ALT levels compared to wildtype mice at 6 hr following APAP overdose^20^, but it is not clear if this can be attributed to loss of PAR1 on hepatocytes or on sinusoidal ECs, where the receptor is most robustly expressed^22^.

To determine the impact of endothelial PAR4 and PAR1 on liver vasculature and hepatocytes after APAP overdose, our current study utilized newly generated *Par4*- and *Par1-* floxed alleles to achieve conditional deletion of the receptors in ECs of adult mice. Our findings highlight: (1) the presence of functional endothelial PAR4 *in vivo,* (2) independent and interdependent roles for PAR1 and PAR4 on liver ECs, and (3) a clarification of the relationship between endothelial PAR-mediated vascular damage and hepatocyte necrosis following APAP overdose.

## MATERIALS AND METHODS

The data that support the findings of this study are available from the corresponding author upon request. Materials and methods can be found in Supplemental Materials, and detailed information about the resources used in this study is listed in the Major Resources Table.

## RESULTS

### Deletion of endothelial *Par1* and/or *Par4* reduces APAP-induced hepatic vascular permeability with distinct kinetics

Endothelial PARs are often studied in the context of vascular permeability, so we initially sought to determine if loss of *Par4* and/or *Par1* on ECs reduced APAP-induced hepatic vascular permeability in mice. We first generated mice with floxed alleles of each receptor. LoxP sites were inserted flanking the first exon of *Par1* and *Par4* (Supplementary Fig. S1A-C), and mice carrying the floxed alleles were identified by PCR genotyping (Supplementary Fig. S1D). In addition to maintaining each floxed line separately, we also generated mice carrying both floxed alleles. This decision was based on the fact that PAR1 and PAR4 can directly interact with each other and can both be activated by thrombin, although PAR1 has a higher affinity for thrombin through its hirudin-like domain (Supplementary Fig. S2)^13^. We next crossed our floxed alleles to the tamoxifen-inducible *Cdh5(PAC)-Cre^ERT2^* line^23^, which is expressed exclusively in ECs when induced in adult mice^24^. The use of an inducible Cre was warranted because the endothelial-specific loss of *Par1* during embryonic development is partially lethal^25^, which would hinder the generation of experimental mice needed for our studies. This approach produced inducible single- and double-EC knockout alleles for *Par1* and *Par4* (hereafter, *Par1^iECko^, Par4^iECko^*, and *Par1/4^iECko^*).

Experimental mice were administered 5 doses of tamoxifen (total of 10 mg) at 5-6 weeks of age for Cre induction. Cre-negative littermate control mice received the same regime. After the final tamoxifen dose, mice were allowed to acclimate for 4 weeks to achieve complete deletion of *Par4* and/or *Par1* and to avoid confounding results from tamoxifen-induced liver dysfunction^26^. Experimental mice were fasted for 16 hr to deplete hepatic glutathione stores and were subsequently administered 300 mg/kg APAP, or a saline vehicle control (Figure 1A). Following the administration of APAP or saline, hepatic vascular permeability was assessed at 6 and 24 hr by assessing leakage of intravascular Evans blue dye (Figure 1B). Basal hepatic permeability at all time points was determined to be 55 ng/mg of Evans blue (shown as a reference line in Figure 1C), with no significant difference between wildtype and *Par^iECko^* mice.

**Figure 1.**
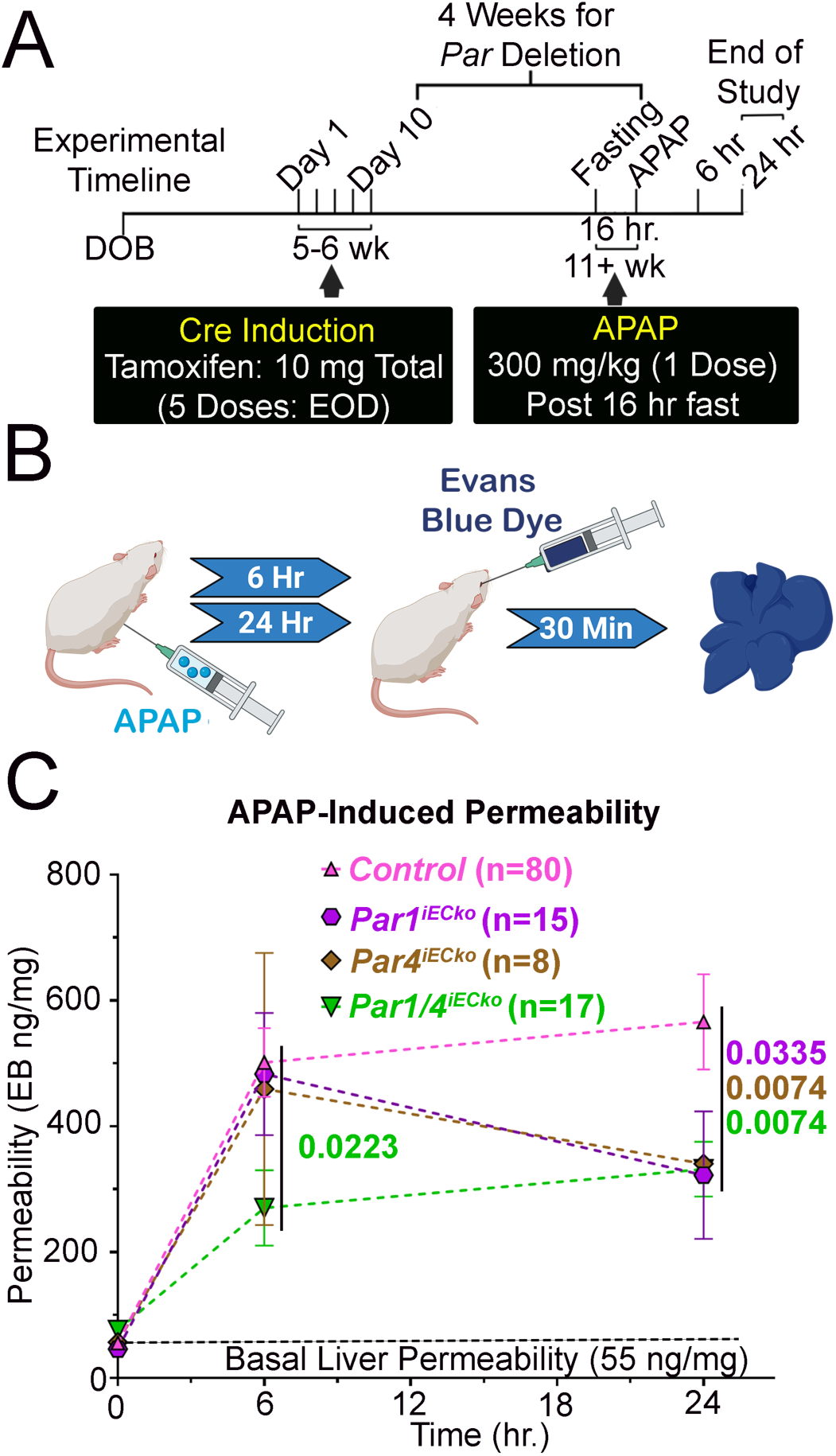
Deletion of endothelial *Par1* and/or *Par4* reduces APAP-induced permeability in mice. (**A**) Outline of experimental procedures performed on mice. DOB: date of birth; EOD: every other day. (**B**) Schematic of Evans blue dye assay for liver permeability. (**C**) Quantification of Evans blue dye extravasated from livers in saline-treated control mice (black dotted line) and APAP-treated control, *Par1^iECko^, Par4^iECko^,* and *Par1/4^iECko^*mice. Data were analyzed by the mixed effects model, and a two-stage linear step-up procedure of Benjamini, Krieger, and Yekutieli was used for multiple comparisons testing. q-values and animal numbers (n) are displayed in the graphs.

Administration of APAP resulted in a significant increase in liver permeability in control mice at 6 hr (∼9.1-fold) and at 24 hr (∼10.2-fold) compared to saline. This APAP-induced vascular permeability was significantly reduced in *Par1/4^iECko^* mice at both 6 hr (∼46.2%) and 24 hr (∼41.3%) compared to Cre-negative littermate controls. By contrast, vascular permeability was not reduced in *Par1^iECko^* mice at 6 hr after APAP overdose but was reduced after 24 hr (∼43.1%). Similarly, vascular permeability was not reduced in *Par4^iECko^* mice at 6 hr after APAP overdose, but it was reduced at 24 hr (∼40.0%), comparable to levels seen in *Par1/4^iECko^* and *Par1^iECko^*mice. These data indicate that loss of endothelial PAR1 and/or PAR4 is sufficient to reduce APAP-induced permeability in the liver, with dual loss of the receptors reducing permeability at 6 hours, and single loss of PAR1 or PAR4 reducing permeability only at 24 hours. These data also demonstrate that PAR4 is a functional receptor on hepatic ECs and that it mediates vascular permeability comparably to PAR1 after APAP overdose.

### Deletion of endothelial *Par1* and/or *Par4* also reduces APAP-induced sinusoidal congestion in mice

Since our data indicated that loss of endothelial PARs reduced APAP-induced vascular permeability, we sought to determine if additional measures of vascular function were rescued in *Par^iECko^* mice. APAP overdose causes congestion of hepatic sinusoids and accumulation of red blood cells (RBCs) around the central veins, overlapping with regions of zone 3 hepatocyte necrosis^27^. This phenotype was observed in our control mice, as evidenced by H&E staining at 24 hr after APAP overdose (Fig. 2). However, we observed reduced RBC congestion in APAP-treated *Par1/4^iECko^* mice compared to littermate controls (∼39.5%; Fig. 2A). We likewise saw reduced congestion in *Par1^iECko^* mice (∼34.6%; Fig. 2B) and *Par4^iECko^*mice (∼35.3%; Fig. 2C) compared to littermate controls. Moreover, *Par1/4^iECko^* did not have significantly more reduction in congestion compared to mice singly deficient in *Par1* or *Par4* (Supplementary Fig. S3A), which suggests that loss of one or both receptors is equally effective at reducing RBC congestion after APAP overdose.

**Figure 2.**
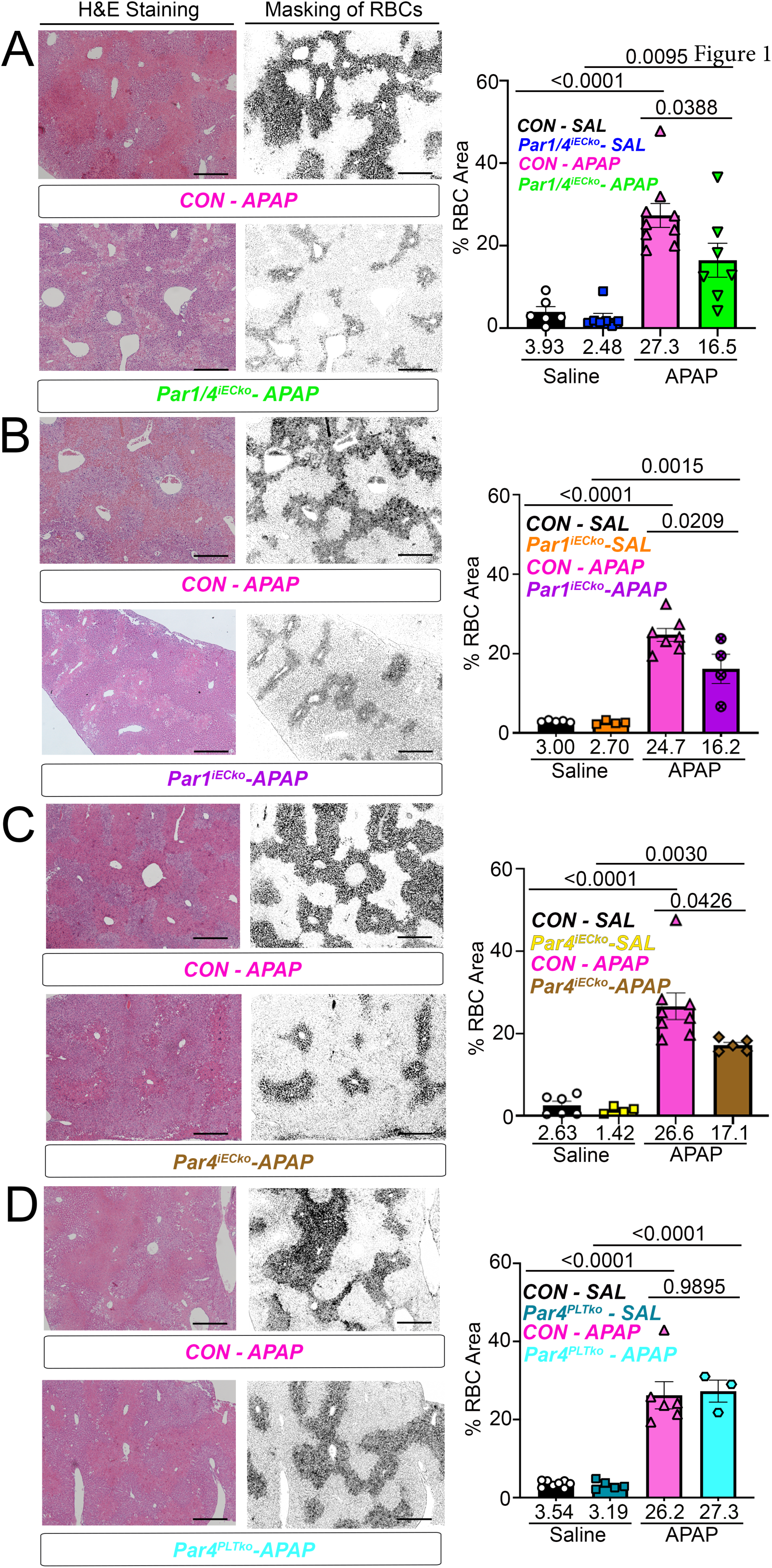
Deletion of endothelial *Par1* and/or *Par4* reduces APAP-induced sinusoidal congestion in mice. (Left) Representative images of H&E staining and masking [to reveal red blood cells (RBCs) in black] in APAP-treated livers from littermate control (CON) versus (**A**) *Par1/4^iECko^*, (**B**) *Par1^iECko^,* (**C**) *Par4^iECko^*, and (**D**) *Par4^PLTko^* mice at 24 hours following overdose. (Right) Quantification of RBC congested areas from histological sections, including in saline (SAL)-treated animals. Data were analyzed by 2-way ANOVA with Tukey’s post hoc test (n=3-9 mice/group). Scale bar = 500 µM.

PAR4 has been best characterized in platelets, where it is highly expressed^22^, and it is typically observed to have low expression levels in ECs^14^. To ensure our rescued RBC congestion phenotype was not influenced by ectopic deletion of *Par4* in platelets, we generated platelet-specific *Par4* knockouts (*Par4^PLTko^*) by crossing *Par4*-flox mice to a *Pf4-Cre* line. Unlike *Par4^iECko^* mice, *Par4^PLTko^* mice did not show reduced RBC congestion compared to littermate controls after APAP overdose (p<0.9954; Fig 2D). Therefore, PAR4 on ECs—rather than on platelets—contributes to sinusoidal congestion after APAP overdose, further confirming that PAR4 is a functional receptor on the hepatic endothelium.

### Deletion of endothelial *Par1* and/or *Par4* does not reduce APAP-induced hepatic injury

To determine if the reduced APAP-induced vascular instability seen in *Par^iECko^* mice correlated with reduced hepatic injury, we collected livers and serum from *Par^iECko^* and littermate controls after APAP overdose. Histological analyses revealed that control mice displayed severe necrosis of zone 3 hepatocytes surrounding hepatic central veins at 6 hr after APAP overdose, and we saw comparable necrosis in livers from *Par1/4^iECko^*, *Par1^iECko^*, *Par4^iECko^*, and *Par4^PLTko^* mice (Fig. 3 and Supplementary Fig. S3B). Since vascular permeability and sinusoidal congestion were most robustly rescued in APAP-treated *Par1/4^iECko^* mice, we further analyzed hepatic injury in these mutants by measuring serum ALT/AST levels at 6 hr and 24 hr after APAP overdose. Serum ALT (Fig. 3E) and AST (Fig. 3F) levels were similar between *Par1/4^iECko^* and littermate controls, indicating that loss of endothelial PARs and reductions in vascular permeability and sinusoidal congestion do not alleviate APAP-induced hepatic injury.

**Figure 3.**
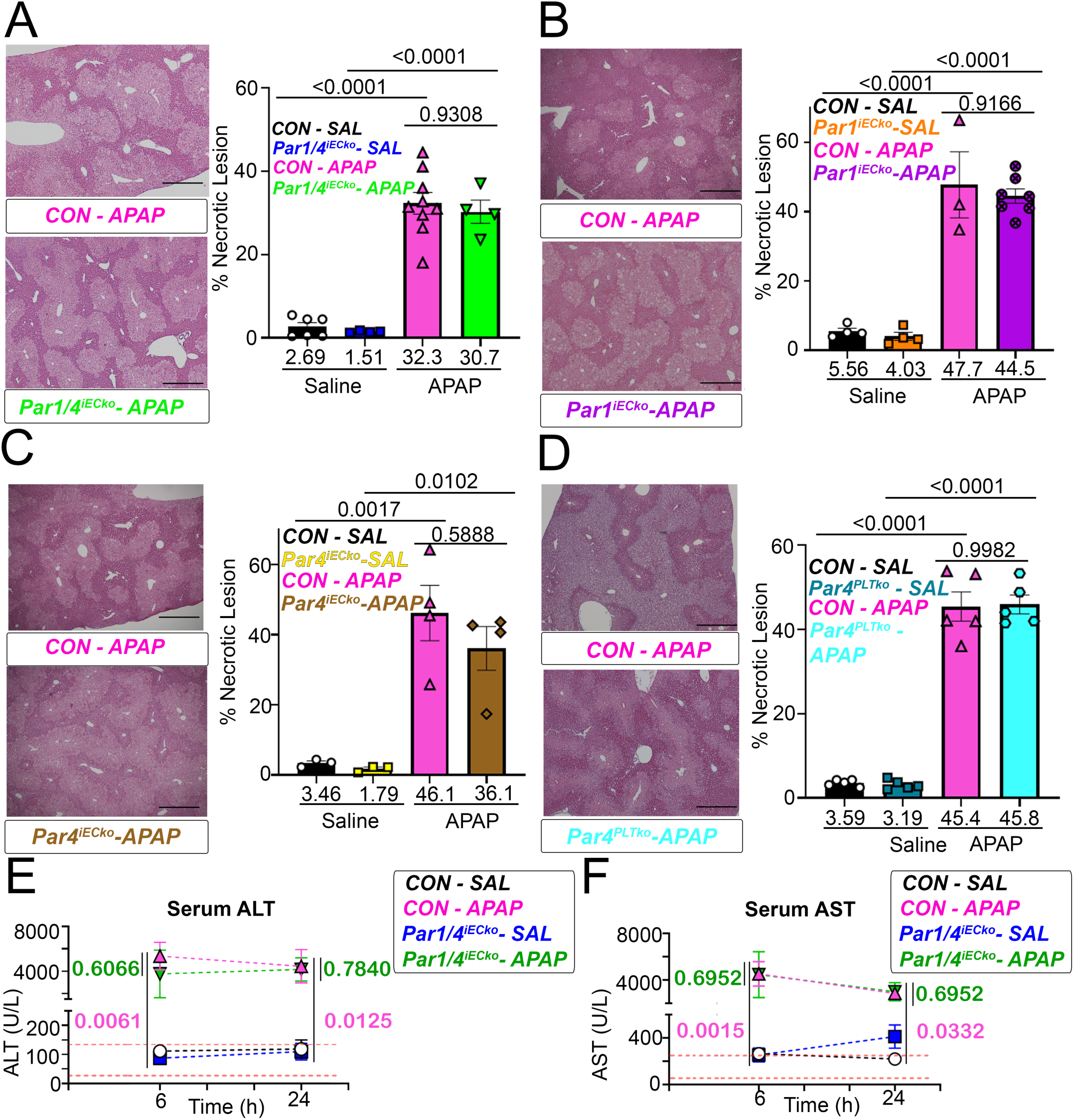
Deletion of endothelial *Par1* and/or *Par4* does not affect APAP-induced hepatocyte necrosis and dysfunction. (**A-D**, Left) Representative images of H&E staining in APAP-treated livers from littermate control (CON) versus (**A**) *Par1/4^iECko^*, (**B**) *Par1^iECko^,* (**C**) *Par4^iECko^*, and (**D**) *Par4^PLTko^* mice at 6 hours following overdose (scale bar = 500 µM). (**A-D**, Right) Quantification of necrotic lesion area from histological sections, including in saline (SAL)-treated animals. (**E-F**) Serum ALT (**E**) and AST (**F**) levels were measured at 6 and 24 hours after saline (SAL) treatment or APAP overdose in littermate control (CON) and *Par1/4^iECko^* mice. The reference range is shown with red dotted lines for ALT (28-132 U/L) and AST (59-247 U/L). (**A-D**) Data were analyzed by 2-way ANOVA with Tukey’s post hoc test (n=3-9 mice/group); p-values are displayed in the graphs. (**E-F**) Data were analyzed by 3-way ANOVA, and a two-stage linear step-up procedure of Benjamini, Krieger, and Yekutieli was used for multiple comparisons testing (n=3-6 mice/group); q-values are displayed in the graphs.

### Translating Ribosome Affinity Purification (TRAP) reveals distinct endothelial responses to APAP overdose

Since endothelial PAR1 and PAR4 contributed to vascular dysfunction following APAP overdose, and GPCR signaling can modulate transcription^28^, we next wanted to assess how PAR1 and PAR4 impacted transcripts in liver ECs using translating ribosome affinity purification followed by next-generation sequencing (TRAPseq; Fig. 4A). We used RiboTag mice for this technique, in which Cre-mediated recombination labels the ribosomal protein RPL22 with an HA tag to facilitate cell-specific isolation of actively translating mRNA. We crossed the RiboTag cassette onto our *Par1^iECko^* and *Par4^iECko^* mice, and we used RiboTag-*Par1^iEChet^* and RiboTag-*Par4^iEChet^* mice as littermate controls since the RiboTag cassette only activates in response to a Cre recombinase. We then collected livers from RiboTag-*Par1^iEChet^*, RiboTag-*Par4^iEChet^*, RiboTag-*Par1^iECko^*, and RiboTag-*Par4^iECko^* mice (3 animals from each genotype) at 6 hr following saline or APAP treatment. Note that the 6 hr time point was chosen for all analyses because it preceded the reduced vascular permeability and sinusoidal congestion seen in *Par1^iECko^* and *Par4^iECko^*mice and might reveal endothelial transcriptional changes associated with those phenotypes. Furthermore, thrombin levels are significantly elevated by this time after APAP overdose^20,21^. Each liver sample was then homogenized and immunoprecipitated to isolate endothelium-specific mRNA from the liver mRNA, thereby allowing us to identify changes in both total liver transcripts and endothelial-specific transcripts after APAP overdose by RNAseq.

**Figure 4.**
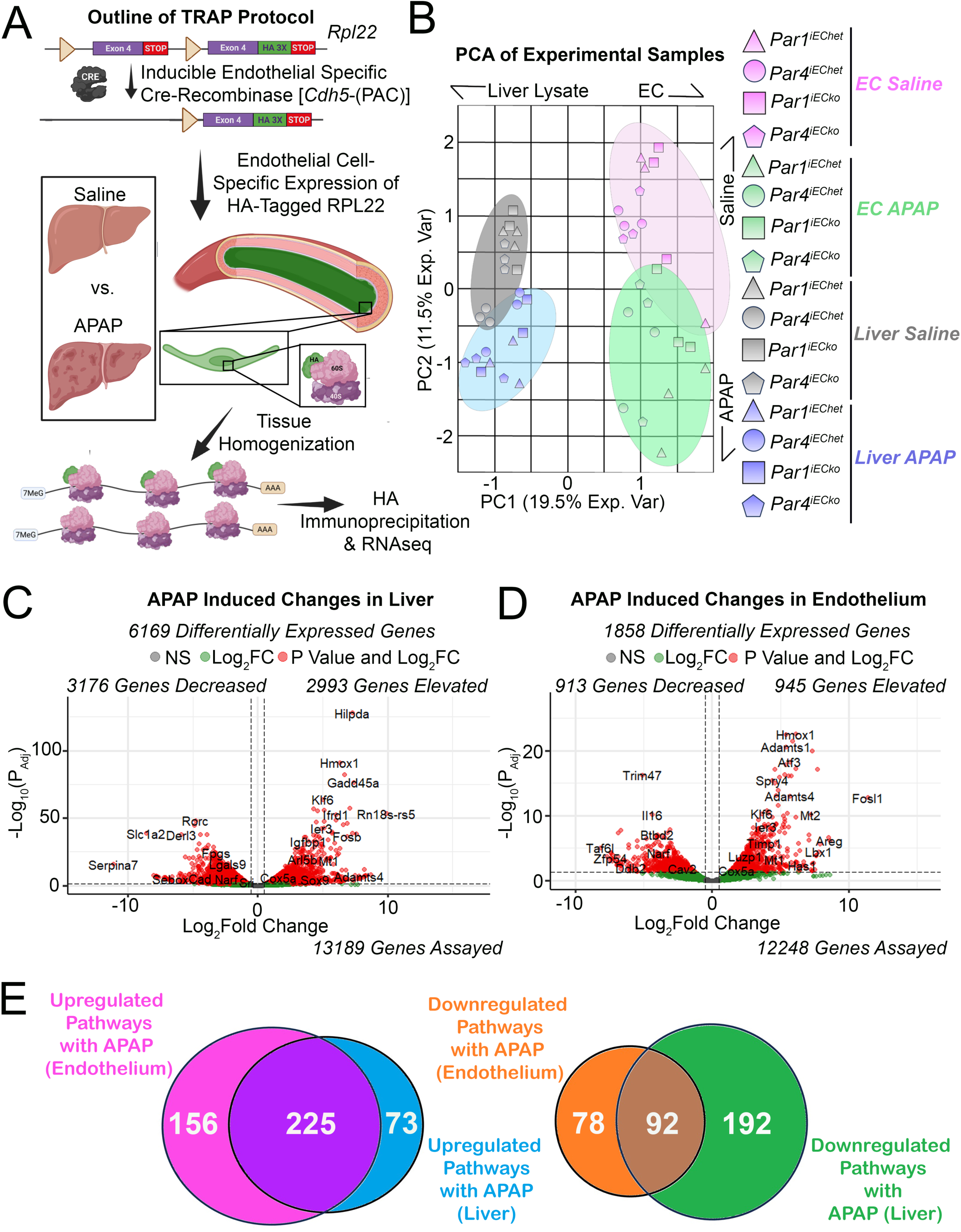
TRAPseq reveals distinct endothelial and total liver responses to APAP overdose. **(A)** Outline of TRAPseq protocol; samples were collected at 6 hr after saline or APAP treatment. **(B)** Principal component analysis (PCA) of RNAseq samples. **(C-D)** Volcano plot of APAP-induced gene expression changes in the liver (C) and endothelium (D). **(E)** Venn diagrams of commonly enriched and depleted pathways between the liver and endothelium after APAP overdose.

Our RNAseq data were then run through a custom analysis pipeline (Supplemental Fig. S4), and the results were subjected to principal component analysis, which showed distinct separation of groups by treatment and fraction (Fig. 4B). In control mice, APAP overdose dramatically affected the transcriptional profiles of the liver when we compared differential gene expression; out of 13,189 genes assayed, 6,169 (∼46.7%) had significant expression changes (P_adj_<0.05) after APAP treatment (Fig. 4C). We next performed Qiagen Ingenuity Pathway Analysis (IPA) on these RNAseq results. Network analysis revealed predicted APAP-induced activation of TNF, IL-1β, IL-6, and EGF (Supplemental Fig. S5A). Furthermore, IPA pathway analysis predicted the upregulation of oxidative phosphorylation, uncoupling proteins, and eukaryotic translation, elongation, and termination in the liver (Supplemental Fig. S5B). These are reasonable predictions since APAP overdose causes oxidative injury and inflammation^19^.

To assess the efficacy of the TRAP enrichment, we compared the transcriptional profiles from endothelium-enriched fractions versus total liver in our saline-treated control animals (Supplemental Fig. S6A). To validate that the TRAP-enriched transcripts were endothelial, we screened the top 500 enriched genes with Gene Ontology (GO; https://geneontology.org/)^29^. The GO terms included pathways commonly associated with ECs, such as “sprouting angiogenesis,” “positive regulation of endothelial cell proliferation,” and “VEGFR signaling pathway” (Supplemental Fig. S6B). We next assessed how APAP overdose affected the endothelial gene expression profiles in control mice. We first removed all TRAP-depleted genes (i.e., genes enriched in non-ECs) from our endothelial dataset. Out of 12,248 genes assayed in the endothelium, we observed significant shifts in 1,858 genes after APAP overdose (∼15.2%; Fig. 4D). IPA network analysis revealed similar driving factors to those we saw in the liver after APAP overdose (i.e., TNF, IL-1β, and EGF) (Supplemental Fig. S5C). However, pathway analysis indicated that endothelial responses predominantly had themes of altered signaling, such as signaling through interleukins, estrogen receptor, phosphatase and tensin homolog (PTEN), and actin-binding rho activating protein (ABRA) (Supplemental Fig. S5D). Comparing alterations in IPA-identified pathways between the liver and endothelium after APAP overdose, we found unique pathways for each fraction (Fig. 4E and Supplemental File 1).

### Endothelial TRAP quantification of *Par1* and *Par4*

Due to the limited expression of PAR4 on only a few cell types^22^, including its low expression in many ECs^30^, single-cell RNAseq (scRNAseq) is usually unable to quantify *Par4* expression since this technique favors the identification of high-expressing transcripts in individual cells^31^. Endothelial TRAP, however, facilitates the aggregation of transcripts from all ECs in an organ, which allowed us to measure actively translating *Par1* and *Par4* transcripts in hepatic ECs. Our TRAP data indicated that both *Par1* and *Par4* are endothelial-enriched genes (Fig. 5). Moreover, using a SALMON RNAseq pipeline, we were able to compare the normalized expression of *Par1* and *Par4* in hepatic ECs; *Par1* was expressed at 85.8 transcripts per million (TPM), and *Par4* was expressed at 0.56 TPM. Therefore, there are 153 *Par1* transcripts for every *Par4* transcript in the hepatic endothelium, whereas there are 43 *Par1* transcripts for every *Par4* transcript in the total liver (Supplemental File 2).

**Figure 5.**
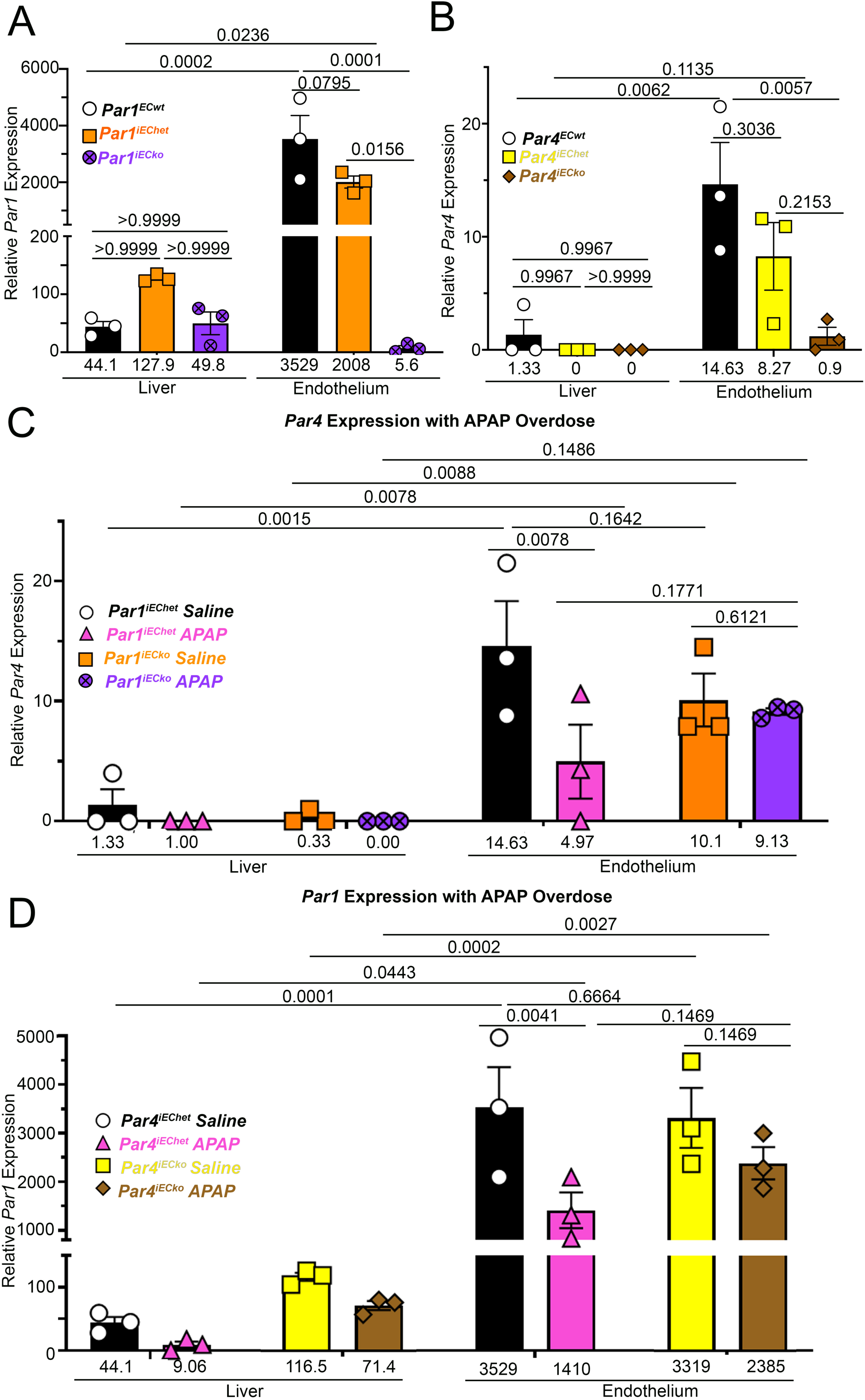
*Par1* and *Par4* expression analysis within TRAPseq data. **(A)** *Par1* expression in liver and endothelium of saline-treated *Par1^ECwt^*, *Par1^iEChet^*, and *Par1^iECko^* mice. **(B)** *Par4* expression in liver and endothelium of saline-treated *Par1^ECwt^*, *Par4^iEChet^*, and *Par4^iECko^* mice. **(C)** *Par4* expression in liver and endothelium from saline- and APAP-treated mice with heterozygous (het) and homozygous (ko) deletion of endothelial *Par1*. **(D)** *Par1* expression in liver and endothelium from saline- and APAP-treated mice with heterozygous (het) and homozygous (ko) deletion of endothelial *Par4*. Data in A and B were analyzed by 2-way ANOVA with Tukey’s post hoc test (n=3); p-values are displayed in the graphs. Data in C and D were analyzed by 3-way ANOVA, and a two-stage linear step-up procedure of Benjamini, Krieger, and Yekutieli was used for multiple comparisons testing (n=3); q-values are displayed in the graphs.

The 153:1 ratio of *Par1* to *Par4* transcripts on hepatic ECs is particularly interesting since our genetic deletion data demonstrate that PAR1 and PAR4 mediate the same vascular responses to APAP overdose (Fig 1C and Supplemental Fig. S3). This indicates that PAR4 is a low-expressing but potent receptor on hepatic ECs. PAR4 undergoes limited desensitization following activation and therefore can strongly amplify signaling through second messengers^12,32,33^, which may account for its robust response in the setting of APAP overdose. Additionally, this disproportionate *Par1:Par4* expression ratio suggests that PAR1-PAR4 heterodimers are unlikely to contribute to a significant portion of the PAR dimer pool on hepatic ECs *in vivo*, especially when compared to PAR1-PAR1 homodimers. Finally, we did not find *Par2* or *Par3* expression in the hepatic endothelium (Supplemental File 2), which makes liver ECs ideal for studying PAR1/PAR4-specific signaling effects in the absence of the other PARs.

### Characterizing how endothelial PARs interact and respond to APAP using TRAPseq

We next sought to understand how the loss of endothelial PAR1 or PAR4 affected transcriptional profiles in the liver and endothelium. We first validated *Par1* and *Par4* deletion from ECs in our RiboTag-*Par1^iECko^*and RiboTag-*Par4^iECko^* mice, respectively. We found an appropriate reduction in *Par1* and *Par4* transcripts in EC fractions from RiboTag-labeled endothelial heterozygous and knockout *Par1* and *Par4* mice, respectively (Fig. 5A, B). *Par1* expression was reduced in the endothelium by ∼99.9% when comparing RiboTag-labeled *Par1^iECko^* and *Par1^ECwt^* mice, while endothelial *Par4* expression was reduced by ∼94% when comparing RiboTag-labeled *Par4^iECko^* and *Par4^ECwt^*mice. We also found that deletion of endothelial *Par4* did not influence *Par1* expression under basal (i.e., saline-treated) conditions, based on the liver and endothelial fractions from RiboTag-labeled *Par4^iEChet^* and *Par4^iECko^* mice (Fig. 5C). Likewise, deletion of endothelial *Par1* did not influence *Par4* expression under basal conditions (Fig. 5D).

Following APAP overdose in RiboTag-*Par1^iEChet^* (control) mice, we found significantly reduced expression of endothelial *Par4* compared to saline-treated control mice (66.0%, p<0.0078; Fig. 5C). However, this reduction in endothelial *Par4* was completely abolished in APAP-versus saline-treated *Par1^iECko^* mice (Fig. 5C), suggesting that PAR1 regulates *Par4* expression after APAP overdose. Similarly, we found reduced expression of endothelial *Par1* (∼61.1%, p<0.0041) in APAP-versus saline-treated RiboTag-*Par4^iEChet^* (control) mice (Fig. 5D). This reduction was diminished in *Par4^iECko^* mice (∼28.1%, p<0.1469) (Fig. 5D), suggesting that PAR4 may likewise regulate *Par1* expression after APAP overdose.

### Endothelial PAR1 and PAR4 differentially regulate hepatic and endothelial gene expression changes to APAP overdose

We next assessed how endothelial *Par1* or *Par4* deletion affects the transcriptional profile of the liver after APAP overdose. First, we compared the liver gene expression profiles between RiboTag-labeled *Par1^iEChet^*mice and *Par1^iECko^* mice at 6 hr after APAP treatment. TRAPseq revealed 345 differentially expressed genes in the *Par1^iECko^* mice (Supplemental Fig. S7A). Subsequent IPA network and pathway analyses indicated a reduction in genes involved in eukaryotic initiation factor 2 (EIF2) signaling (Supplemental Fig. S7B and C). Notably, when all the pathways that were altered by APAP overdose in the livers of RiboTag-*Par1^iEChet^* mice were considered, only a small fraction (∼7%) were rescued in RiboTag-*Par1^iECko^* mice (Supplemental Fig. S7D). We also compared the liver gene expression profiles from RiboTag-labeled *Par4^iEChet^* mice and *Par4^iECko^* mice after APAP overdose, but we did not find any significant differences between these control and mutant samples (Supplemental Figure S8).

Finally, we sought to identify how endothelial deficiency of PAR1 and PAR4 affected hepatic endothelial gene expression after APAP overdose. We first compared the endothelium of RiboTag-labeled *Par1^iEChet^* mice and *Par1^iECko^* mice after APAP overdose. TRAPseq revealed 49 differentially expressed genes (Fig. 6A), and IPA pathway analyses indicated a reduction in genes involved in the pathways of “Rho GTPase cycle,” and “phagosome formation” (Fig. 6C). IPA network analysis of overlapping pathways from these samples suggested the “Rho GTPase cycle” to be a central node (Supplemental Fig. S9). Further analysis revealed an increase in the expression of several genes involved in processes related to cytoskeletal rearrangement, phagocytosis, and vascular permeability in APAP-treated control endothelium, and these genes were rescued in APAP-treated *Par1^iECko^* endothelium (Fig 6D). Alterations in genes involved in the Rho GTPase cycle and cytoskeletal rearrangement are notable since PAR1 is known to regulate these processes post-translationally^34,35^. These new TRAP data indicate that PAR1 may also impact these cellular processes at the level of transcription. We also found upregulated expression of AP-2 complex subunit beta (*Ap2b1*) in APAP-treated control endothelium and rescue of this gene in APAP-treated *Par1^iECko^*endothelium (Fig 6D). Since AP-2 is involved in the endocytosis and trafficking of PAR1 and PAR4^36,37^, *Ap2b1* transcriptional upregulation following PAR1 activation may facilitate increased trafficking of PARs to maintain EC thrombin responsiveness after APAP overdose. Note that downregulated *Dll4* served as a positive control for our dataset since it has been reported to drop in expression in ECs after APAP overdose (Fig 6D)^38^. Finally, when we compared the endothelium of RiboTag-labeled *Par4^iEChet^* mice and *Par4^iECko^* mice after APAP overdose, we did not find significant differences between the samples (Fig. 6B). Therefore, PAR1 mediates transcriptional changes in ECs with APAP overdose, but PAR4 does not.

**Figure 6.**
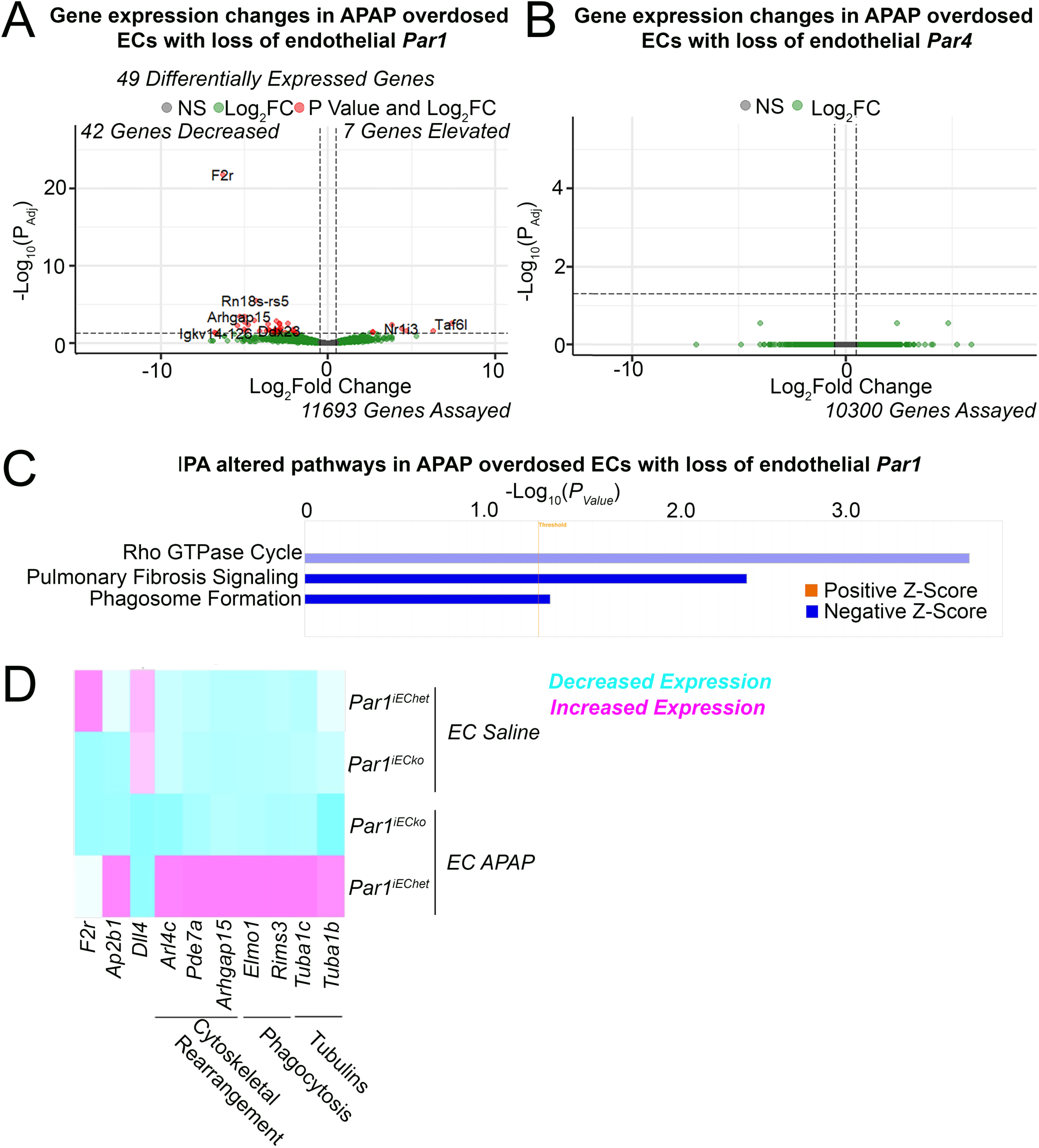
TRAPseq reveals that endothelial PAR1 and PAR4 differentially regulate EC gene expression changes in response to APAP overdose. **(A-B)** Volcano plots of gene expression changes between control and *Par1^iECko^* ECs (A) or *Par4^iECko^* ECs (B) after APAP overdose. **(C)** IPA pathways that were differentially regulated between ECs from control and *Par1^iECko^* mice after APAP overdose. **(D)** Heatmap of select genes differentially expressed in ECs between control and *Par1^iECko^* mice after saline treatment or APAP overdose.

## DISCUSSION

After two decades of controversy, our study demonstrates the presence of functional PAR4 on hepatic ECs *in vivo*. Our TRAP data reveal that both *Par1* and *Par4* are enriched in ECs of the murine liver, although endothelial *Par4* transcripts are expressed at low levels compared to *Par1* (1:153; Supplemental File 2). Importantly, *Par4^iECko^* mice phenocopy both the reduced sinusoidal congestion and vascular permeability phenotypes seen in *Par1^iECko^* mice after APAP overdose, indicating that endothelial PAR4 is a potent receptor despite its low expression.

Our study also shows that endothelial PAR1 and PAR4 contribute to hepatic vascular permeability and sinusoidal congestion after APAP overdose, since these phenotypes were attenuated in *Par^iECko^* mice. Interestingly, dual loss of endothelial *Par1/4* reduced hepatic permeability at both early (6 hr) and late (24 hr) time points after APAP overdose, while loss of either individual receptor only reduced permeability at 24 hr, suggesting that endothelial PAR1 and PAR4 act redundantly to promote vascular instability 6 hours after APAP overdose.

Another important insight from this study is that vascular permeability does not have a significant impact on the hepatocyte necrosis seen at 6 hr after APAP overdose. Centrilobular hepatocyte necrosis is driven by elevated production of N-acetyl-p-benzoquinone imine (NAPQI), a toxic intermediate of APAP metabolism that is generated by the enzyme CYP-4502E1, which is primarily expressed in centrilobular hepatocytes^39,40^. Because of the hepatocyte-specific expression of CYP-4502E1, it is reasonable that NAPQI production and subsequent hepatocyte necrosis would not be altered in endothelial *Par*-deficient mice after APAP overdose. We believe previous studies that used global KO strategies to deplete PARs likely saw reduced hepatocyte necrosis after APAP overdose due to the loss of low-expressing PARs on hepatocytes^20,21^. Future studies can use hepatocyte-specific deletion of PARs to test this hypothesis.

Concordant changes in both hepatic vascular permeability and sinusoidal congestion after APAP overdose are understandable since both are measures of vascular stability. Previous studies have shown that sinusoidal ECs display gaps after APAP overdose through which RBCs can extravasate^27,41–43^. We observed comparably reduced congestion/bleeding in both single- and double-*Par1/4^iECko^* mice at 24 hr after APAP overdose. PAR1 can regulate endothelial shape change and vascular permeability either through Gα_q_/phospholipase C-β /calmodulin/myosin light chain kinase signaling or through cytoskeleton rearrangements mediated by Gα_12/13_/Rho-GTPase signaling^44^. PAR4 also regulates actin rearrangements in ECs^45^. Deletion of endothelial PARs likely makes liver ECs functionally insensitive to elevated thrombin levels, which are associated with APAP overdose^20,21^. As a result, reduced thrombin-induced EC shape changes may account for increased vascular stability in *Par^iECko^*mice.

Although our study addresses the existence and function of PAR4 on liver ECs, we recognize that some of the initial controversy about endothelial PAR4 arose because of conflicting data generated from ECs in other organs. For example, Kataoka et al demonstrated that PAR4 is functional in murine pulmonary ECs in 2003^15^, but others have since failed to reproduce those results^4^. This made us curious about *Par4* expression levels in ECs from lungs and other organs. Therefore, we surveyed *Par1* and *Par4* expression in murine ECs from multiple organs by looking at datasets generated by Kalucka et al^30^ and Cleuren et al^17^. We found that *Par1* was expressed in ECs of most organs that were analyzed. *Par4*, on the other hand, had endothelial expression limited to the kidney, heart, liver, colon, and spleen (Supplemental Figure S10). This suggests that endothelial PAR4 may function most prominently in these organs *in vivo* and is consistent with the roles we identified for PAR4 in hepatic ECs in this study.

We also surveyed an RNAseq dataset by Buglak et al^46^ and found that cultured HUVECs exposed to shear stress undergo significant upregulation of *PAR4* (∼13.3-fold), although similar upregulation is not seen for *PAR1* (Supplemental Figure S11). Concordantly, scRNAseq analysis of human liver^47^ indicates that endothelial *PAR4* expression is most prominent around the portal vein, where flow rates are highest^48^ (Supplemental Figure S12D). Likewise, increased PAR4 levels have been detected in higher caliber pulmonary vessels, which have higher flow rates^49^. Since PAR4 is activated by circulating proteases, and high blood flow rates would decrease the local ligand residency time and availability for a given receptor, these expression data suggest that high protease concentrations would be required for PAR4 activation under high flow conditions.

Altogether, PAR4 appears to be a selectively expressed and low-affinity thrombin receptor on ECs in contrast to PAR1. However, our study indicates that PAR4 is a potent receptor that is capable of mediating similar responses to PAR1 in hepatic ECs. We hypothesize that the primary role of endothelial PAR4 is to maintain thrombin responsiveness in scenarios where the enzyme is present at high concentrations and has saturated endothelial PAR1 signaling. In addition, because PAR4 has limited desensitization mechanisms and can strongly amplify second messengers^12,32,33^, its low expression is not indicative of a failure to function, but rather, it serves to limit the signaling of the receptor except in response to high thrombin activity, such as occurs with APAP overdose.

Notably, our TRAPseq data demonstrate that PAR1 and PAR4 show distinct differences in influencing EC transcriptional reprogramming after APAP overdose. Two interesting genes were *Par1* and *Par4* themselves, especially since the receptors facilitated downregulation of each other after APAP overdose. We attribute these changes to be due to receptor trans-downregulation, in which the activation of one PAR downregulates the other. We identified 49 other targets that were differentially expressed in *Par1*-deficient ECs, but no other genes were affected by *Par4* deletion. Given that both *Par1^iECko^*and *Par4^iECko^* mice display a reduction in permeability and sinusoidal congestion after APAP overdose, the gene expression changes in *Par1^iECko^* mice are unlikely to result from secondary effects of stabilizing the hepatic vasculature but rather from direct loss of PAR1 in hepatic ECs.

Several of the genes that we found differentially expressed in *Par1^iECko^*mice are linked to cellular permeability. These gene changes may not be functionally consequential in APAP overdose, since both *Par1^iECko^*and *Par4^iECko^* mice had reduced vascular permeability after APAP overdose even without comparable transcriptional changes, but they may be relevant in other models of thrombin-induced permeability or in organs lacking endothelial PAR4 (Supplemental Figure S10). Given that endothelial PAR4 is a potent signaling receptor and lacks desensitization mechanisms like PAR1, it might be best to limit PAR4 signaling effects to post-translational activities since unregulated transcription could be deleterious to long-term EC health and function. Altogether, our findings indicate that endothelial PAR1 and PAR4 are functionally redundant in regulating vascular stability but are distinct in their expression, signaling, and potency.

## LIMITATIONS OF STUDY

TRAPseq represents a leap forward in our ability to quantify low-expressing genes like *Par4*. Although this technology allowed us to identify the ratio of actively translating *Par1:Par4* transcripts in hepatic ECs *in vivo*, one limitation we have is that we cannot claim that this ratio reflects the ratio of the receptors at the EC surface. Future studies using genetic mouse models to label endogenous proteins *in vivo*, such as FUNCAT^50^, may be able to provide a better approximation of the true *in vivo* EC surface expression ratio of these receptors. Another limitation to this study is the lack of mechanistic insight into how PARs reduce endothelial permeability and sinusoidal congestion in the liver. Theoretically, we might have used cultured ECs to address this question, but this study highlights how static ECs are unsuitable for studying PAR4. Moreover, other labs have shown that PAR4 can mediate permeability by regulating cytoskeleton rearrangement and EC shape change through G protein-dependent and -independent signaling^45,51,52^. Therefore, we elected to focus instead on addressing the long-standing controversy on whether PAR4 is even present and functional in the vascular endothelium through our in vivo studies.

## ACKNOWLEDGEMENTS

We thank Jun Xie, M.D., for assistance with mouse colony maintenance. We thank Christopher Bottoms, Ph.D., for his help with the OMRF cluster. Data processing and analysis were supported by the OMRF Center for Biomedical Data Sciences. Diagrams were created with BioRender.com.

## SOURCES OF FUNDING

This work was supported by a grant to C.T.G. from the National Institutes of Health (R35HL144605) and predoctoral fellowships to R.R. from the American Heart Association (23PRE1014240) and OMRF.

## AUTHOR CONTRIBUTIONS

R.R. designed and conducted all experiments and analyses, including the writing of all the informatics programs used in the manuscript. A.C.A.C. prepared immunoprecipitated RNA into libraries for sequencing. R.R. wrote the manuscript. C.T.G. oversaw the project and edited the manuscript. All authors provided critical feedback that helped shape the research, analysis, and manuscript.

## DATA AVAILABILITY

The data reported in this paper have been deposited in the Gene Expression Omnibus (accession no. GSE266395).

## NON-STANDARD ABBREVIATIONS AND ACRONYMS

ALT: Alanine Aminotransferase
APAP: Acetaminophen
AST: Aspartate Aminotransferase
ECs: Endothelial Cells
GO: Gene Ontology
GPCR: G Protein-Coupled Receptor
HUVECs: Human Umbilical Vein Endothelial Cells
IPA: Ingenuity Pathway Analysis
NAPQI: N-acetyl-p-benzoquinone imine
PAR1: Protease-Activated Receptor 1
PAR4: Protease-Activated Receptor 4
RBCs: Red Blood Cells
TPM: Transcripts Per Million
TRAP: Translating Ribosome Affinity Purification

## DISCLOSURES

The authors have no conflicts of interest to disclose.

## HIGHLIGHTS

- PAR4 is a low-expressing but highly potent receptor on hepatic endothelial cells *in vivo*.
- PAR1 and PAR4 can act synergistically on hepatic endothelial cells but also possess distinct functions following APAP overdose.
- Alterations in hepatic vascular stability do not contribute to hepatocyte dysfunction after APAP overdose.
- The utilization of translating ribosome affinity purification (TRAP) combined with conditional genetic knockouts for GPCRs represents an innovative *in vivo* method for studying receptor-mediated transcriptional reprogramming following a pathological insult like APAP overdose.

## Notes

### Competing Interest Statement

The authors have declared no competing interest.

### Summary of Updates

Supplemental Materials have been added, including Supplemental Figures S1-S12, Methods and Materials, Supplemental Tables ST1-ST3, Major Resource Table ST4, Supplemental References

