## Supplemental Materials for "Acetaminophen Overdose Reveals Protease-Activated Receptor 4 as a Low-Expressing but Potent Receptor on the Hepatic Endothelium"

**Supplemental Figures**

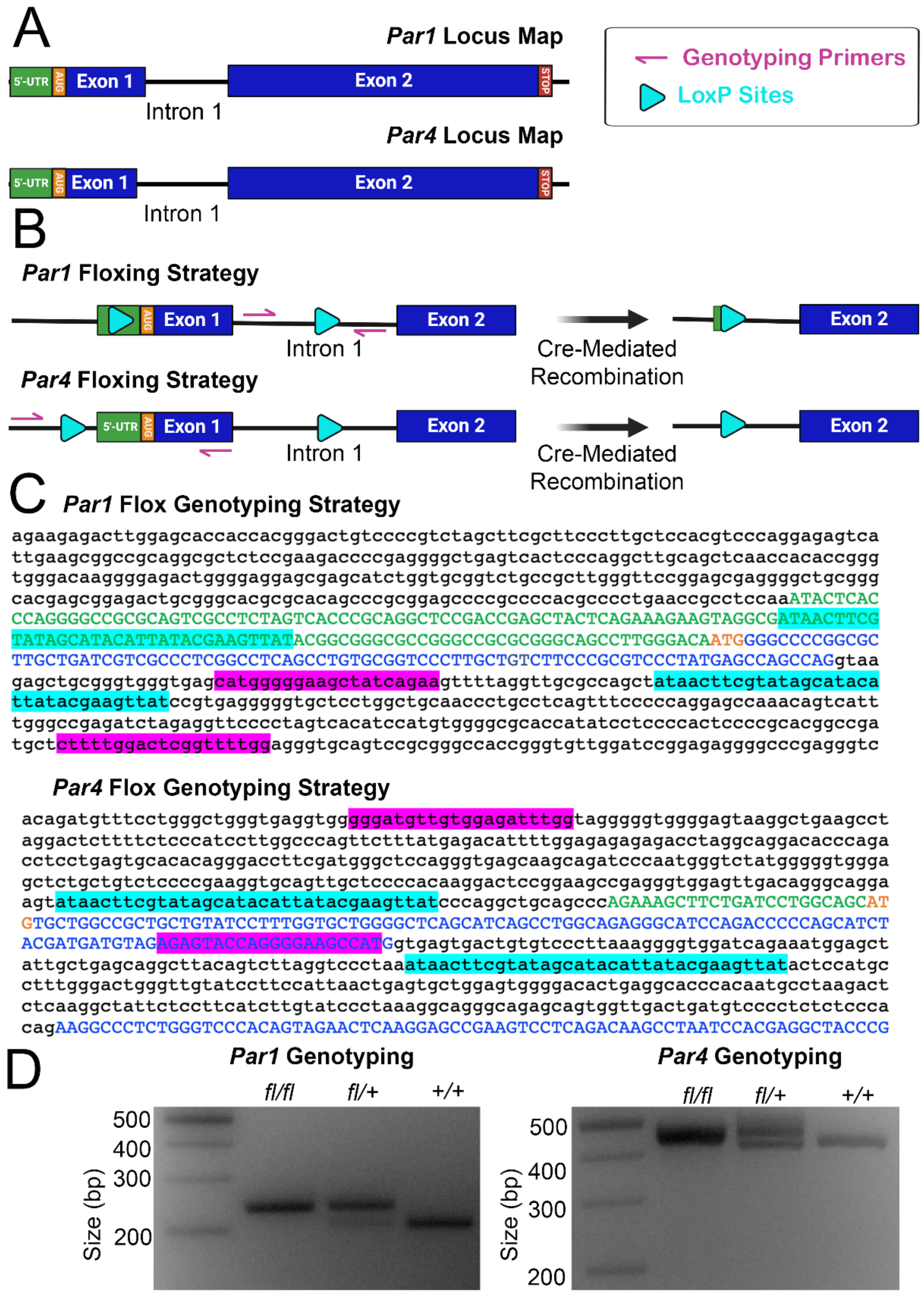

**Supplemental Fig. S1: Mouse *Par1* and *Par4* locus maps and floxing strategy. (A)** Maps of the murine *Par1* and *Par4* loci. Both genes are composed of two exons separated by one intron; exon 1 is smaller than exon 2 for both genes. **(B)** Floxing strategies for the *Par1* and *Par4* genes. For the *Par1* locus, LoxP sites (cyan) were inserted in the 5'-untranslated region [5'-UTR **(Green)**] and intron 1, thus flanking exon 1 **(blue)**. Genotyping primers (pink) were designed to flank the 3' LoxP site. For the *Par4* locus, LoxP sites were inserted upstream of the 5'-UTR and in intron 1, thus flanking exon 1. Genotyping primers were designed to flank the 5' LoxP site. For both alleles, Cre-mediated recombination results in the deletion of exon 1. **(C)** Sequencing results of the modified *Par1* and *Par4* loci with highlighted LoxP sites and genotyping primers. **(D)** DNA gels showing PCR-based genotyping reactions from mice with modified *Par1* and *Par4* alleles.

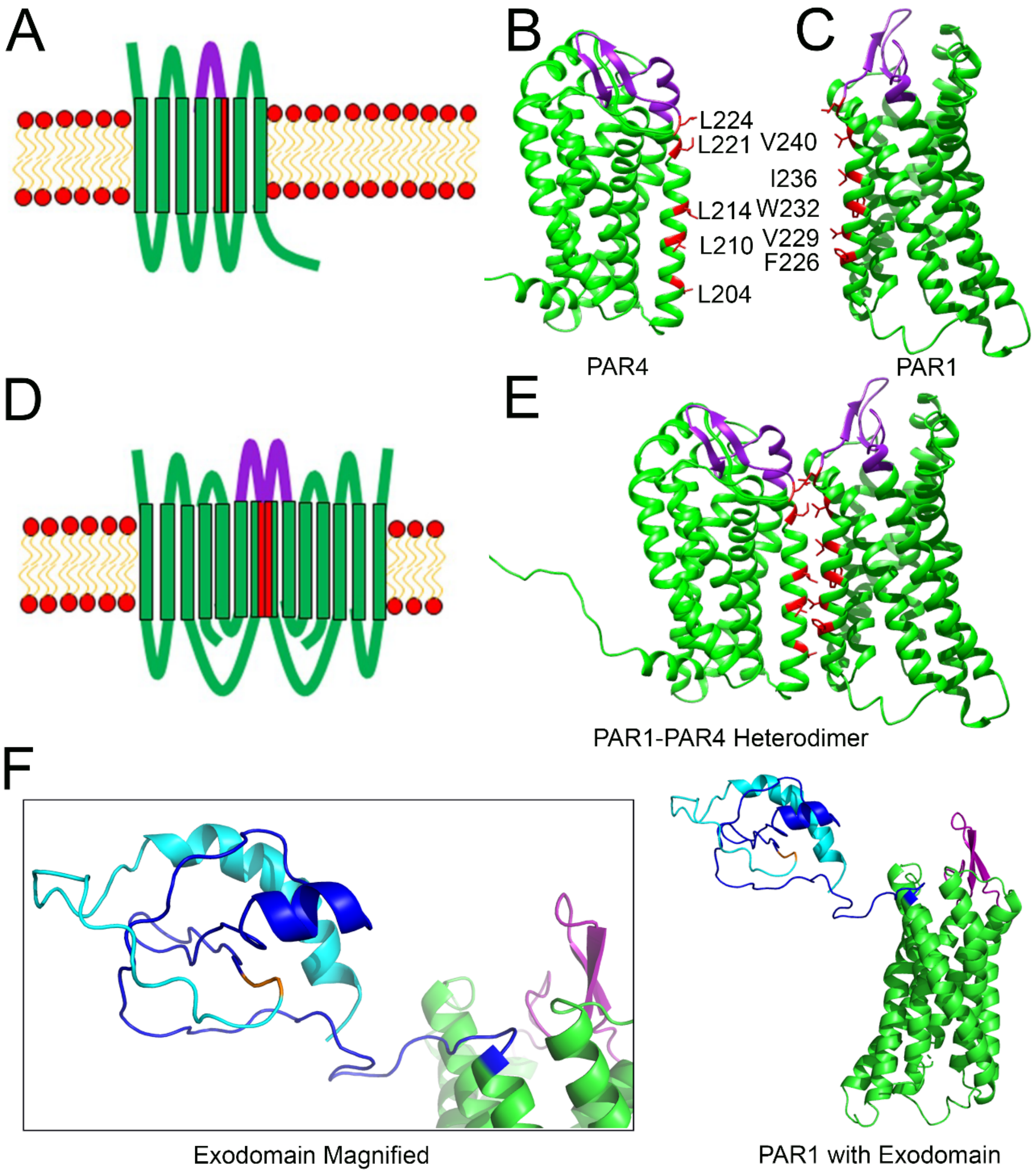

**Supplemental Fig. S2: Murine PAR1 and PAR4 structures. (A)** Schematic of a PAR in the plasma membrane with the extracellular domain 2 (EC2) shown in **purple** and the transmembrane 4 domain (TM4) domain with putative dimerizing residues shown in **red**. **(B)** *In silico* model of murine PAR4 with EC2 domain shown in **purple** and putative dimerizing residues in **red**. **(C)** *In silico* model of murine PAR1 with EC2 domain shown in **purple** and putative dimerizing residues in **red**. **(D)** Schematic of dimerized PARs with binding between both TM4 domains. **(E)** In *silico* models of murine PAR1 and PAR4 rearranged to show dimerization along TM4. (**F**) *In silico* model of murine PAR1 and the N-terminal exodomain with a hirudin-like site that potentiates thrombin binding. The cleaved peptide is shown in **cyan**, the cleavage site is shown in **orange**, and the tethered ligand is shown in **blue**, which includes the hirudin-like site (**blue α-helix**). The EC2 is shown in **purple**.

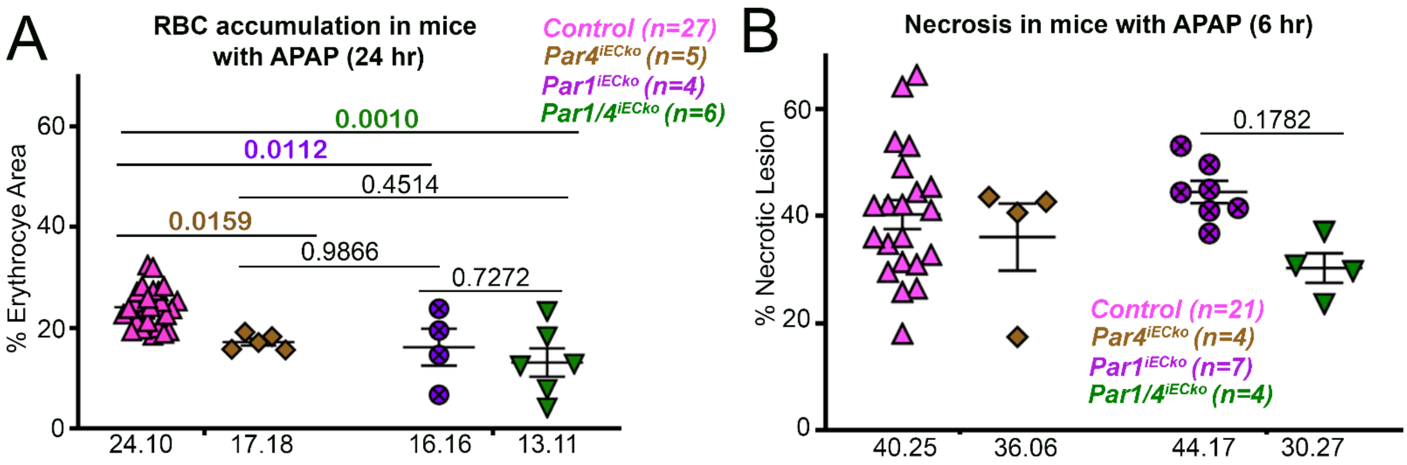

**Supplemental Fig. S3: Deletion of endothelial *Par1* and *Par4* reduces APAP-induced hepatic sinusoidal congestion but not hepatocyte necrosis. (A)** Quantification and comparison of hepatic red blood cell (RBC) areas between all genotypes of mice at 24 hr after APAP overdose. Outlier analysis was performed (Rout Q=1% performed and outliers were removed: 3 values from control mice and 1 value from *Par1/4$^{iECko}$* mice); the remaining data were analyzed by 2-way ANOVA with a Tukey's post hoc test (n=4-30 mice/group). **(B)** Quantification and comparison of necrotic lesions between all genotypes of mice at 24 hr after APAP overdose. Data were analyzed by 2-way ANOVA with a Tukey's post hoc test (n=4-21 mice/group).

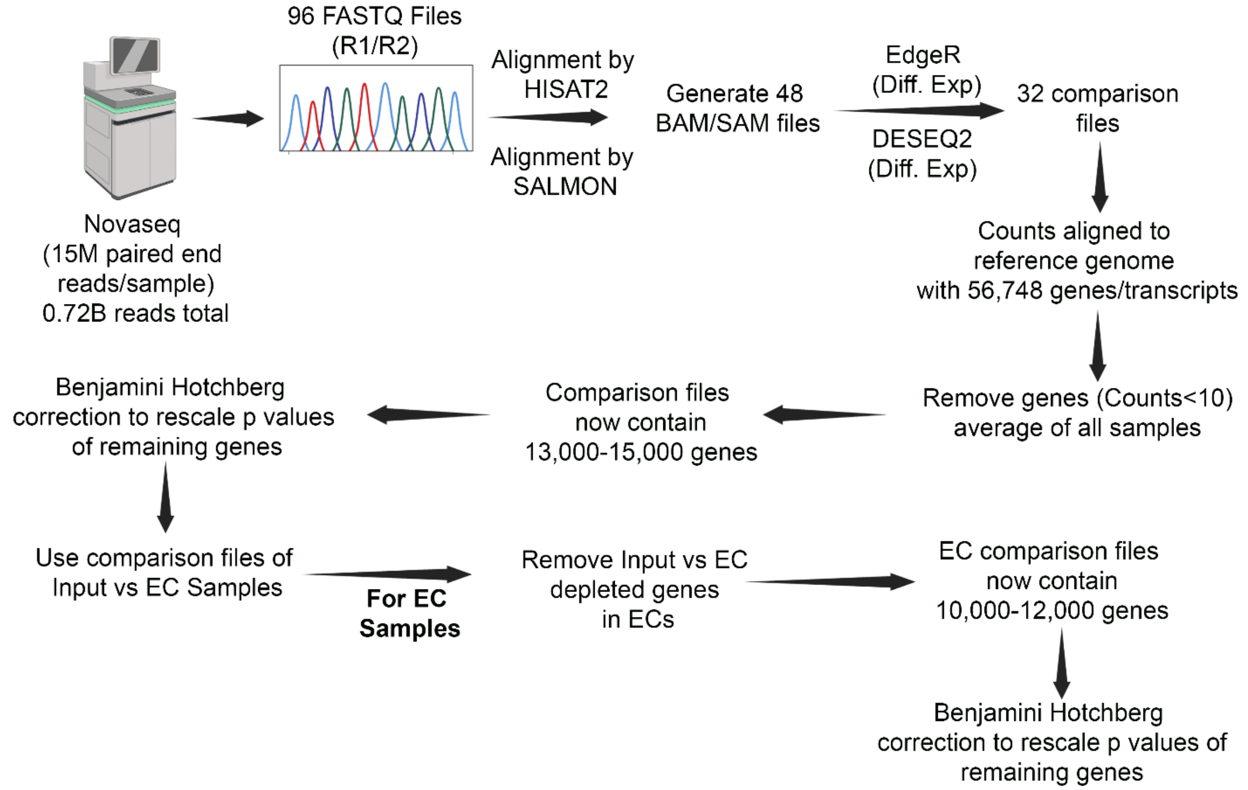

**Supplemental Fig. S4: Workflow for RNAseq data analysis.** Schematic of RNAseq analysis from FASTQ file processing to final generation of count matrices.

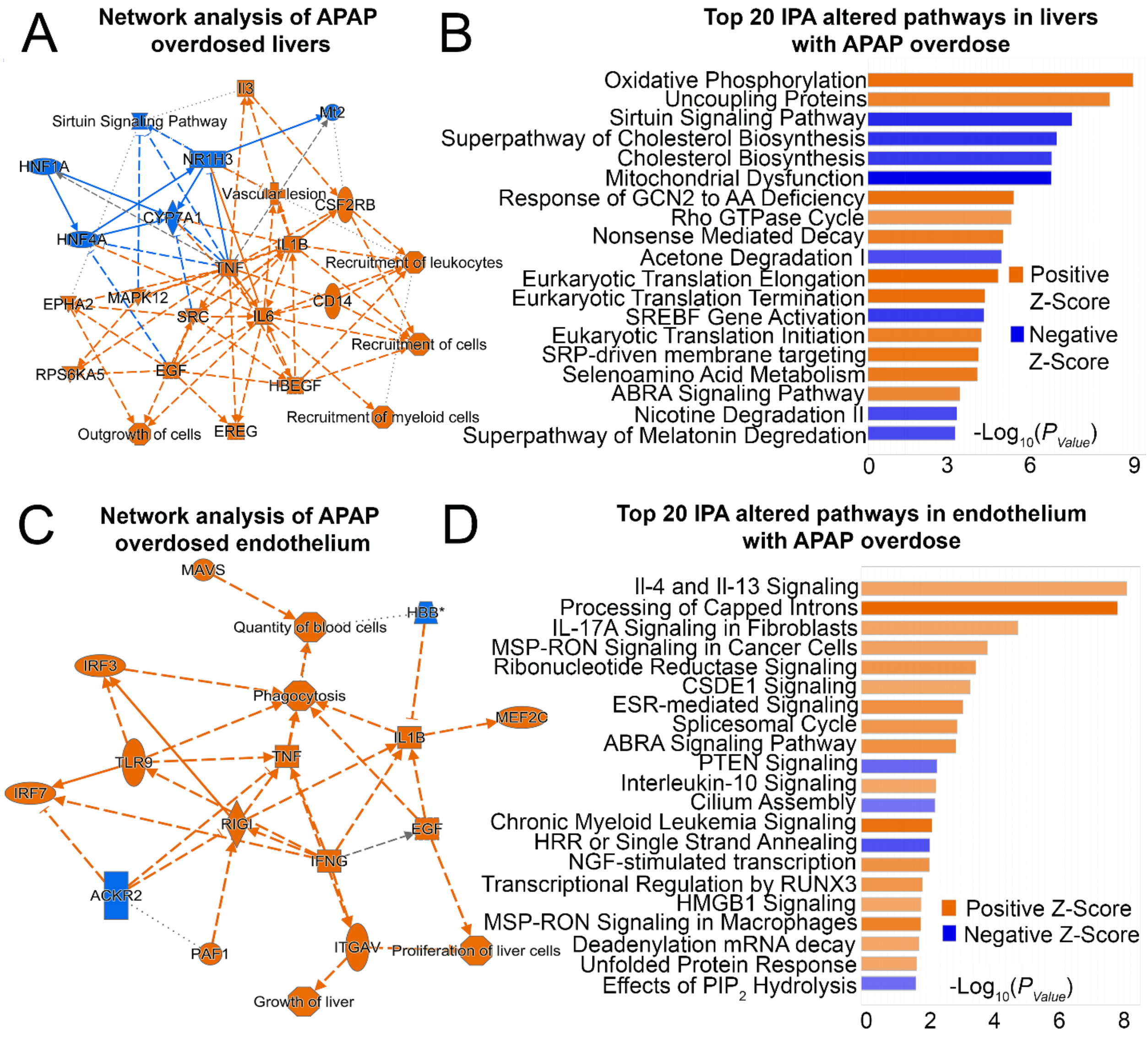

**Supplemental Fig. S5: IPA analysis of liver and endothelium at 6 hr after APAP overdose. (A)** Qiagen IPA network analysis of livers after APAP overdose. **(B)** Top 20 altered pathways in livers after APAP overdose **(C)** Qiagen IPA network analysis of the endothelium after APAP overdose. **(D)** Top 20 altered pathways in the endothelium after APAP overdose. For A and B, 6,169 significantly differentially expressed genes were identified in livers after APAP overdose (see Fig. 5C); genes with PAdj < 0.05, $Log_2FC > ABS|2|$, and FDR < 0.01 were selected for IPA analysis, yielding a subset of 2,089 genes. For C and D, 1,858 significantly differentially expressed genes were identified in the endothelium after APAP overdose (see Fig. 5C), and 1,844 of those genes were selected for IPA analysis.

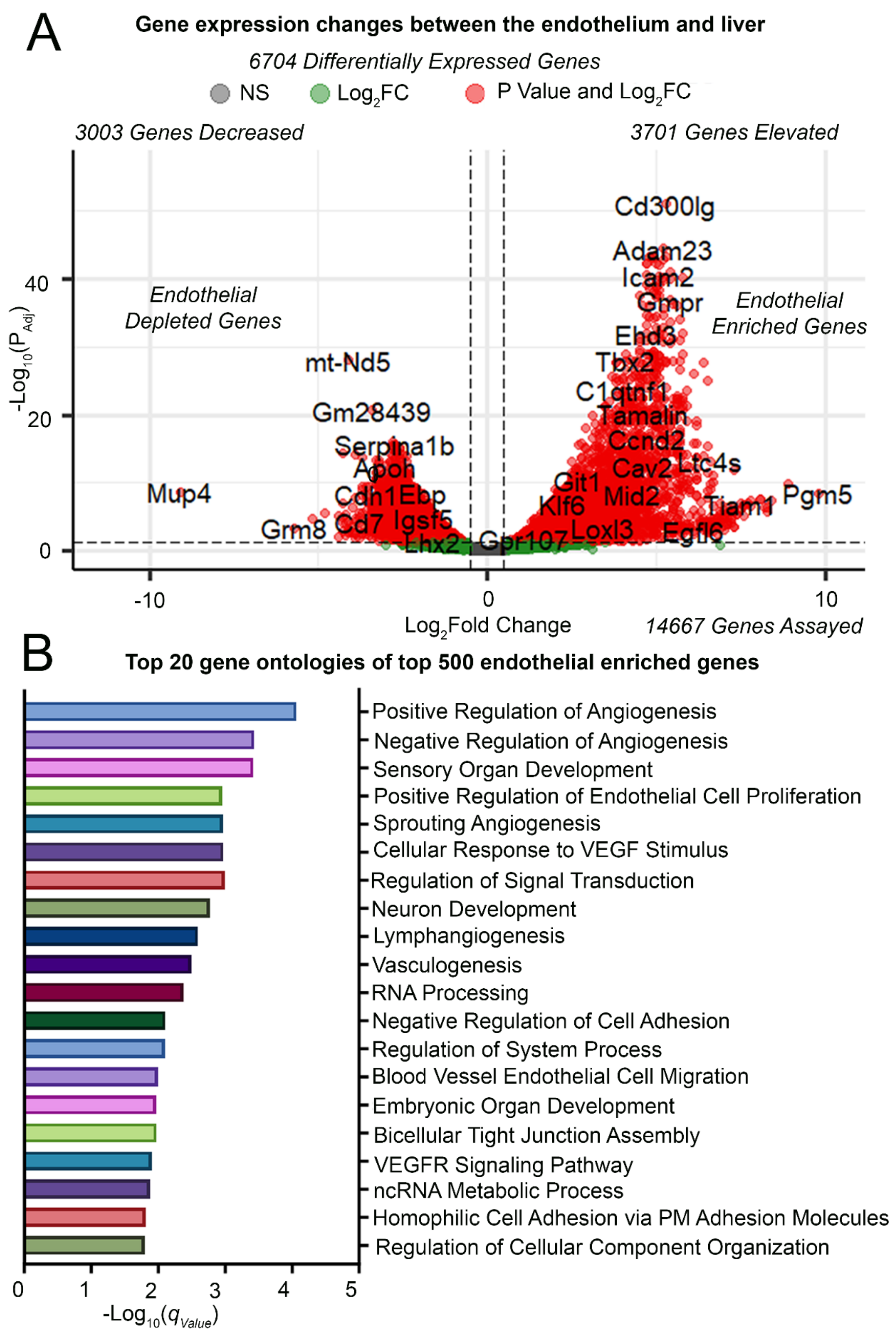

**Supplemental Fig. S6: TRAP enrichment of transcripts in the endothelium. (A)** Volcano plot of enriched and depleted genes when comparing the endothelium fraction to the total liver. **(B)** Top 20 Gene Ontologies from the top 500 enriched genes found in A.

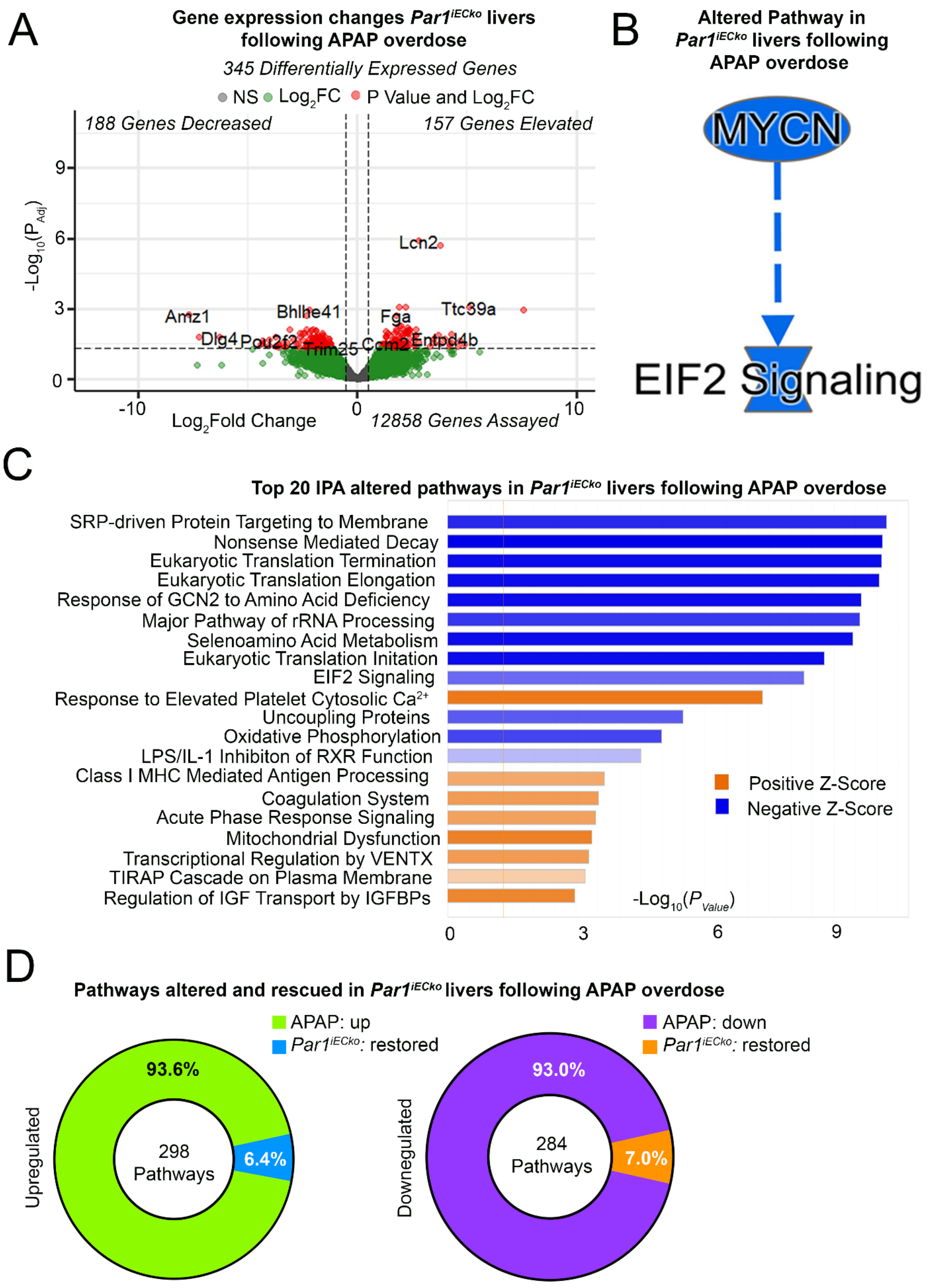

**Supplemental Fig. S7: Deletion of endothelial *Par1* marginally alters liver transcriptional profiles at 6 hr after APAP overdose.** **(A)** Volcano plot of differentially expressed genes between control and *Par1$^{iECko}$* livers after APAP overdose. **(B)** Qiagen IPA network analysis between control and *Par1$^{iECko}$* livers after APAP overdose. **(C)** Top 20 altered pathways between control and *Par1$^{iECko}$* livers after APAP overdose. **(D)** The proportion of up- and downregulated pathways in livers that retain or change their gene expression profile with loss of endothelial *Par1*. ‘Restored’ designates pathways in the *Par1*$^{iECko}$ group that show expression levels restored to those seen in saline-treated control livers.

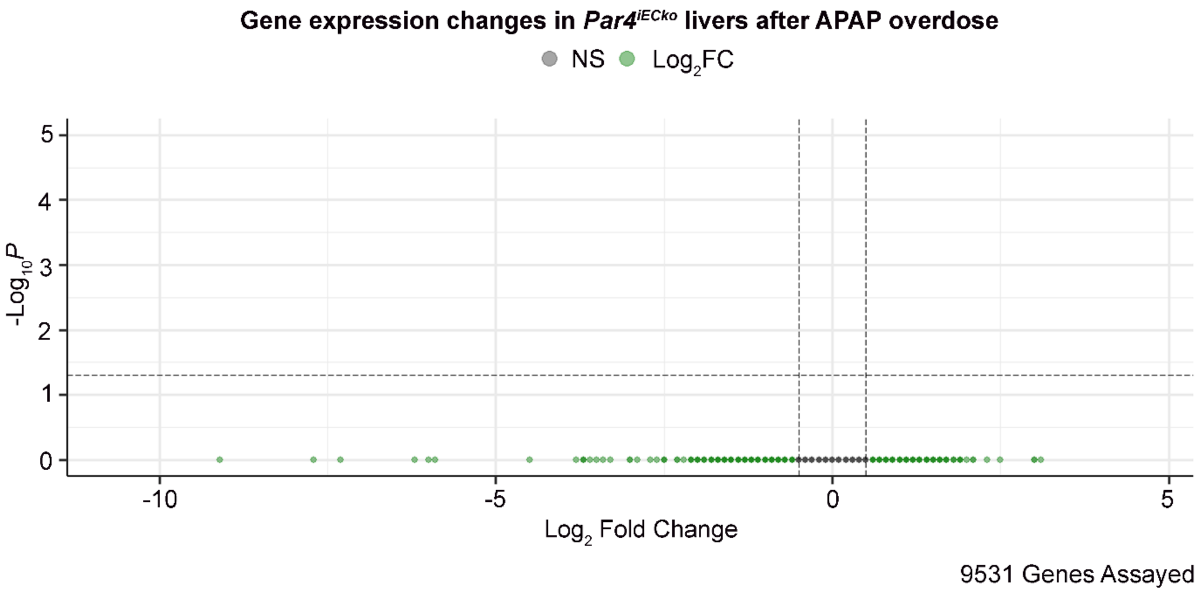

**Supplemental Fig. S8: Transcriptional profiles are not altered in *Par4[iECko]* livers at 6 hr after APAP overdose.** Volcano plot of differentially expressed genes between control and *Par4[iECko]* livers after APAP overdose.

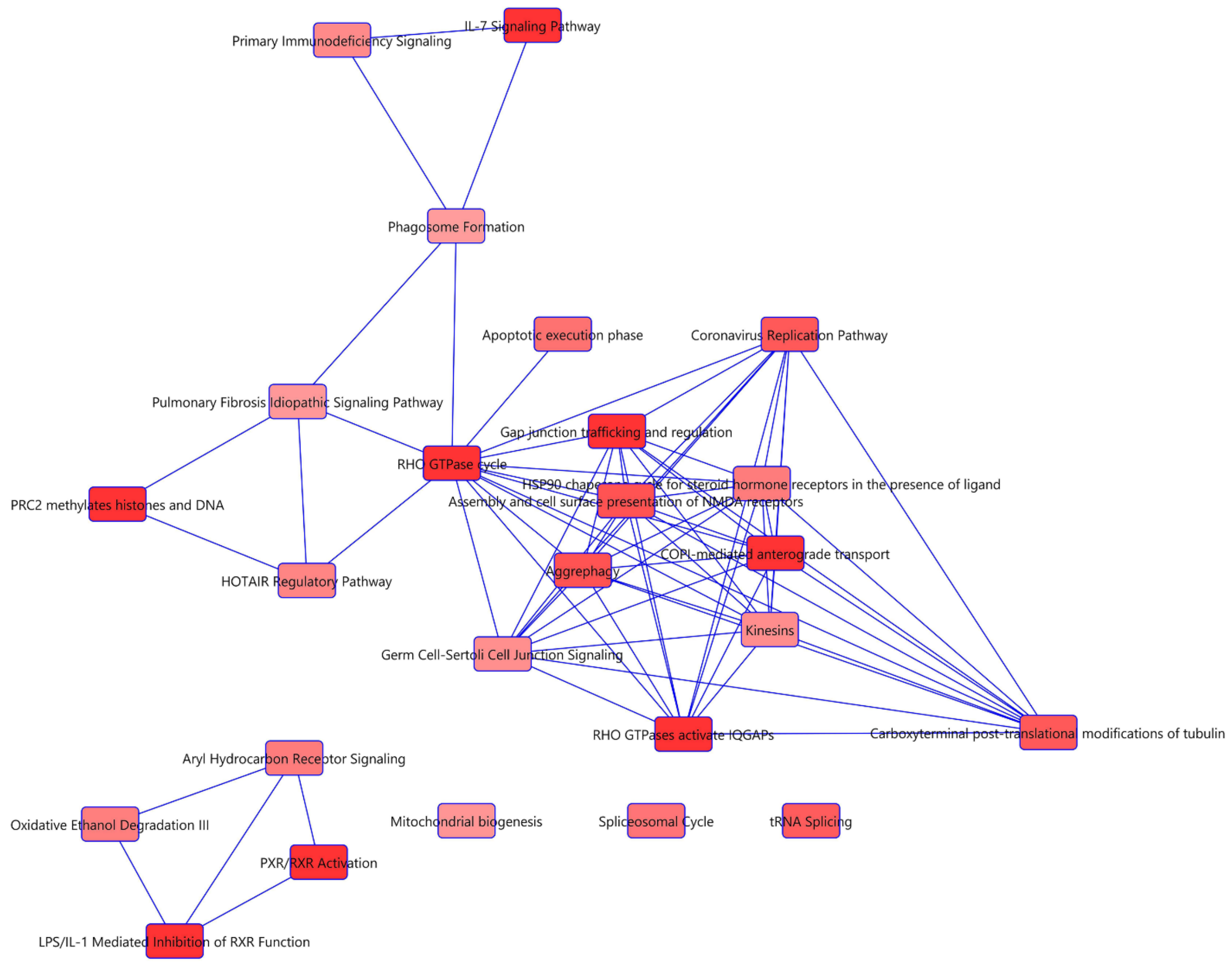

**Supplemental Fig. S9: IPA overlapping pathway analysis of *Par1[iECko]* endothelium at 6 hr after APAP overdose.** IPA network map of shared and overlapping pathways altered in *Par1[iECko]* endothelium after APAP overdose.

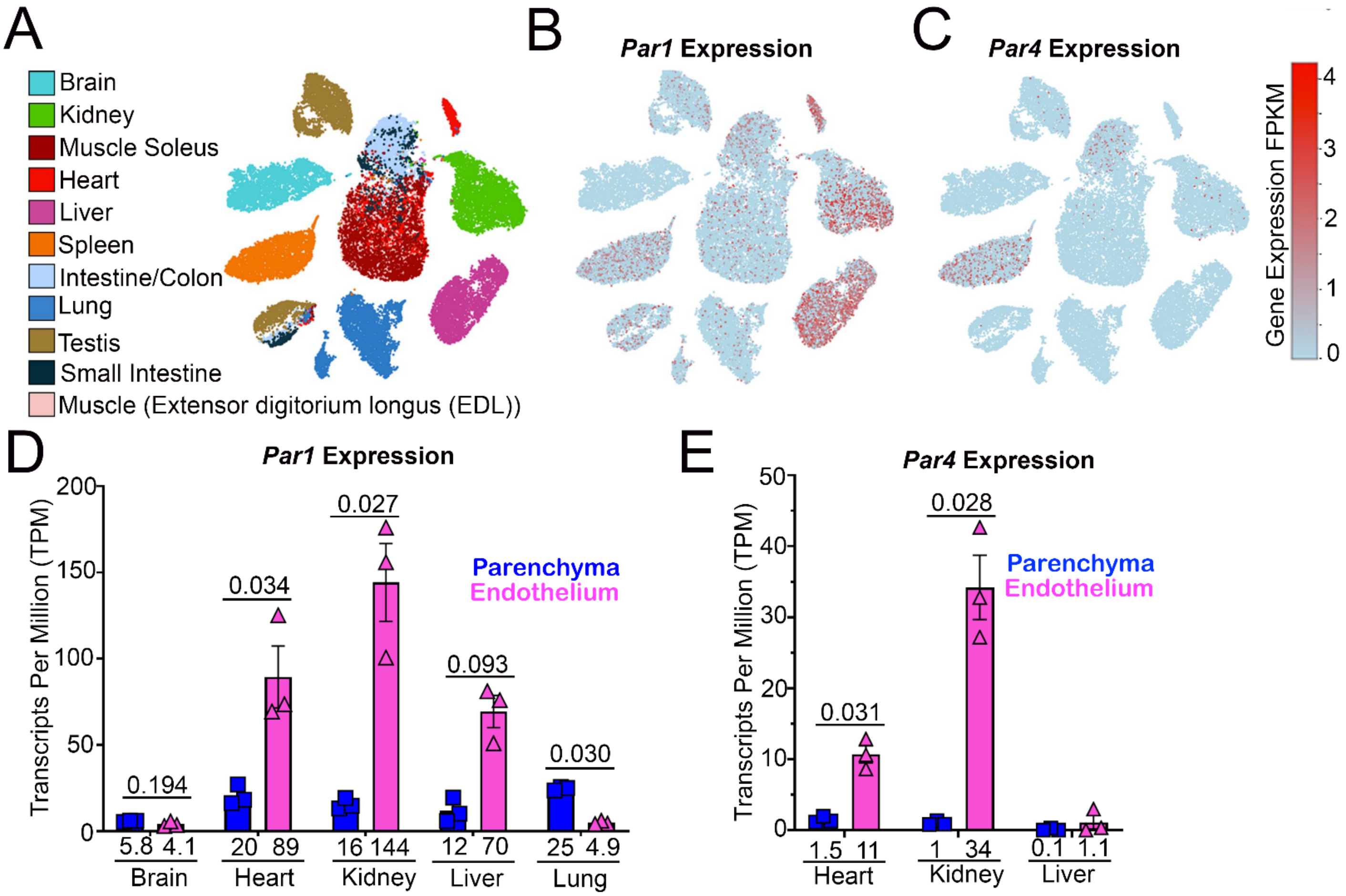

**Supplemental Fig. S10: Organotypic endothelial *Par1* and *Par4* expression. (A-C):** Analysis of scRNAseq data generated by Kalucka et al[1]. **(A)** UMAP of scRNAseq data showing ECs originating from different murine tissue beds. **(B)** *Par1* expression in fragments per kilobase of transcript per million mapped reads (FPKM) between ECs from different murine tissue beds. **(C)** *Par4* expression in FPKM between ECs from different murine tissue beds. **(D-E)**: Analysis of TRAP data generated by Cleuren et al[2]. **(D)** TPM counts of *Par1* expression in lysates and ECs of different murine tissue beds. **(E)** TPM counts of *Par4* expression in lysates and ECs of different murine tissue beds. Data in D and E were analyzed using Welch's t-test with a Benjamini-Hochberg correction, as performed in the original publication; q-values are displayed in the graphs.

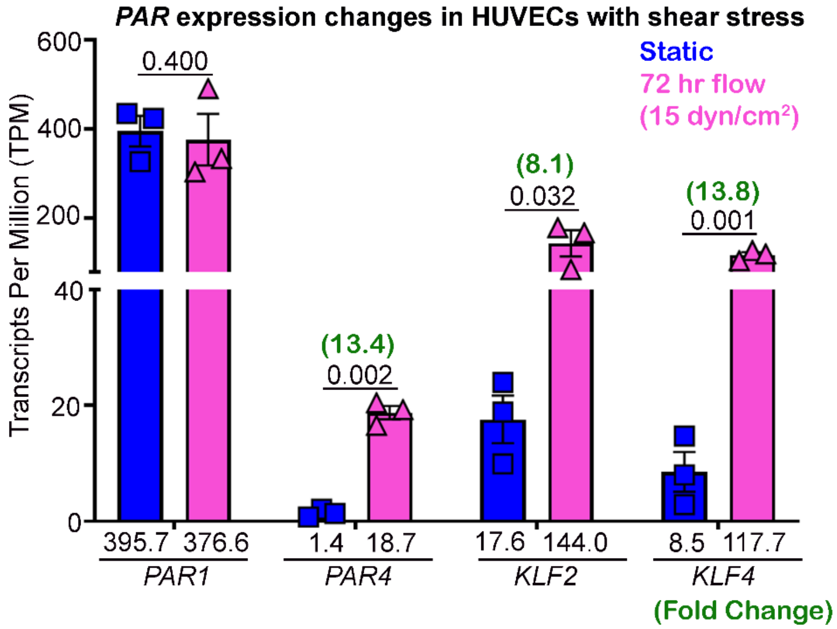

**Supplemental Fig. S11: *PAR1* and *PAR4* expression in HUVECs with shear stress.** *PAR1* and *PAR4* expression (TPM counts) in primary human umbilical vein ECs (HUVECs) subjected to shear stress (15 dyn/cm$^2$) for 72 hr compared to cells cultured in static conditions; data were extracted from Buglak et al[3]. *KLF2* and *KLF4* are shown as examples of classical flow-sensitive genes. Fold change calculations are shown in green. Data were analyzed using a Welch's t-test after validation of data normality by a Shapiro-Wilk normality test; multiple comparisons were performed by setting the desired false discovery rate (FDR) to 1% and performing a two-stage step-up method of Benjamini, Krieger, and Yekuteli; q-values are displayed in the graph.

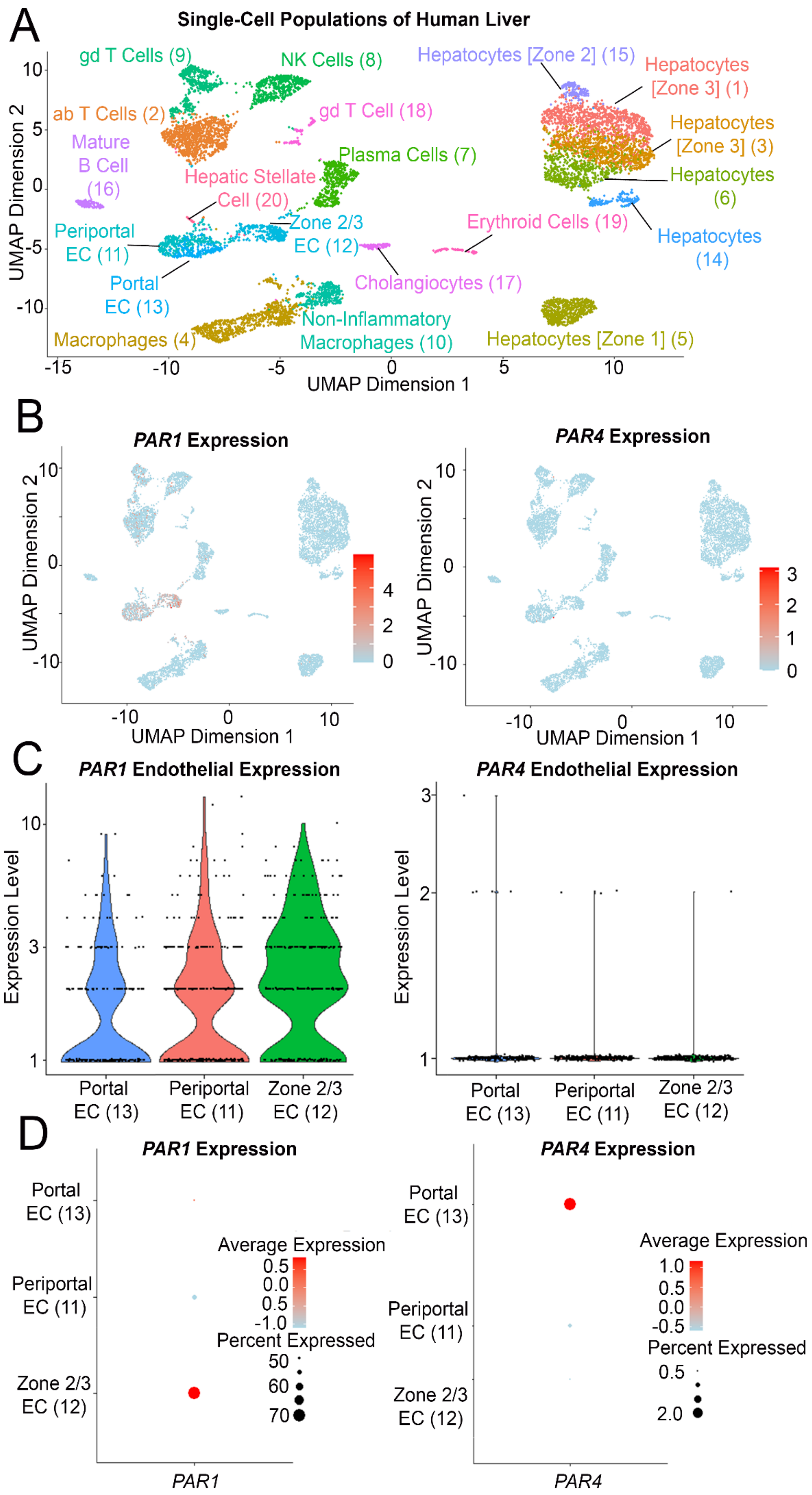
A
Single-Cell Populations of Human Liver
gd T Cells (9)
NK Cells (8)
Hepatocytes [Zone 2] (15)
Hepatocytes [Zone 3] (1)
ab T Cells (2)
gd T Cell (18)
Hepatocytes [Zone 3] (3)
Mature B Cell (16)
Plasma Cells (7)
Hepatic Stellate Cell (20)
Hepatocytes (6)
Periportal EC (11)
Zone 2/3 EC (12)
Erythroid Cells (19)
Hepatocytes (14)
Portal EC (13)
Cholangiocytes (17)
Non-Inflammatory Macrophages (10)
Macrophages (4)
Hepatocytes [Zone 1] (5)
UMAP Dimension 1
UMAP Dimension 2
B
PAR1 Expression
PAR4 Expression
C
PAR1 Endothelial Expression
PAR4 Endothelial Expression
Expression Level
Portal EC (13)
Periportal EC (11)
Zone 2/3 EC (12)
D
PAR1 Expression
PAR4 Expression
Average Expression
Percent Expressed
PAR1
PAR4

**Supplemental Fig. S12: scRNAseq of the human liver shows *Par1* and *Par4* expression in ECs of distinct zones. (A-D)** Analysis of scRNAseq data generated by MacParland et al[4]. **(A)** UMAP of scRNAseq data showing 20 distinct cell types in the human liver. **(B)** *PAR1* and *PAR4* expression in FPKM between different liver cells. **(C)** Violin plots and **(D)** dot plots of *PAR1* and *PAR4* expression in portal, periportal, and zone 2/3 ECs.

#### Methods and Materials

**Mice.** Mice used in this study include:

- *Par1*-flox (Generated for our lab by ViewSolid Biotech, Inc.)
- *Par4*-flox (Generated for our lab by ViewSolid Biotech, Inc.)
- *Cdh5(PAC)-Cre*$^{ERT2}$ (gift of Ralf Adams, Max Planck Institute for Molecular Biomedicine; available through Taconic: #13073)[5]
- *Pf4-Cre* (The Jackson Laboratory: #008535)[6]
- *Rpl22*$^{tm.1.1Psam}$ (RiboTag; The Jackson Laboratory: #029977)[7]

Inducible EC-specific deletion of *Par1* and/or *Par4* was achieved by crossing mice expressing *Cdh5(PAC)-Cre*$^{ERT2}$ with *Par1*-flox and/or *Par4*-flox mice. Platelet-specific deletion of *Par4* was achieved by crossing mice expressing *Pf4-Cre* with *Par4*-flox mice. For TRAPseq experiments, the RiboTag allele was crossed onto *Par1*$^{iECko}$ and *Par4*$^{iECko}$ lines; RiboTag-*Par1*$^{iEChet}$ and RiboTag-*Par4*$^{iEChet}$ mice were used as controls for comparison. Genotyping of mutant alleles was conducted with the primers listed in Supplemental Table 1. All mice were maintained on a C57Bl/6J background under environmental conditions that were previously described[8]. Gene deletion with the *Cdh5(PAC)-Cre*$^{ERT2}$ line was performed in 5–6-week-old mice with the administration of five doses of 10 mg/mL tamoxifen (200µL) (Sigma: #T5648) dissolved in peanut oil via oral gavage every other day, as described[9]. Mice were then housed for 4 weeks before further experimentation at 11 weeks of age or older (average weight of 25 g). All experimental animal protocols were approved by the Institutional Animal Care and Use Committee at the Oklahoma Medical Research Foundation. All animal studies were done in adherence with the ARRIVE guidelines proposed by the National Centre for the Replacement Refinement and Reduction of Animals in Research (NC3Rs). Guidelines include: designing, conducting, interpreting, and reporting animal studies.

**Generation of *Par1*- and *Par4*-flox mice.** Deletion of the first exon of the *Par1* gene results in partial embryonic lethality, thus demonstrating the essential role of this exon[10]. Therefore, we generated an allele in which the first coding exon (residues 1-29) was flanked by loxP sites, using CRISPR/Cas9, using a similar genetic strategy to the PAR1 global knockout mice[11]. Deletion of the exon results in loss of the translational start codon (ATG) and the signal peptide with subsequent loss of protein translation. A CRISPR/Cas9 gRNA was synthesized with specificity for the *Par1* locus, and a repair template for homology-directed repair. The homology arms were selected based on the gRNA cleavage sites, then assembled with two loxP sites on both sides of exon 1 for *Par1.* Cas9 mRNA, a *Par1*-specific gRNA, and a donor DNA repair template were co-microinjected into murine zygotes during the active DNA replication period. 10 to 14 days after birth, pups were genotyped by PCR to identify the founders (F0) with the desired gene modification. The founder mice were analyzed by sequencing at the targeted locus to confirm the genotyping results. Since mutations made by the CRISPR/Cas9 system may be mosaic in founder mice, selected founders were bred with wildtype mice to obtain F1 heterozygotes, whose alleles were confirmed by PCR genotyping. *Par4,* which has an identical makeup of exons and introns to *Par1*, was floxed in the same manner.

**Acetaminophen overdose.** Acetaminophen (APAP; Sigma A5000) was dissolved in 0.9% saline at a concentration of 15 mg/mL. To mitigate APAP oxidation and degradation, the drug was stored in paraffin-wrapped tubes and protected from light. Sealed and wrapped aliquots of APAP were kept under constant vacuum pressure in a desiccator. Starting at 11+ weeks of age, mice were fasted overnight (~16 hr) and then injected intraperitoneally with 300 mg/kg APAP. Food was returned to animal cages immediately after the APAP injection. Animals were euthanized at 6 hours or 24 hours after APAP administration to collect tissue and blood for analysis since these times correlate with two types of liver pathology: centrilobular hepatocyte necrosis presenting at 6 hours (1), and centrilobular bleeding presenting at 24 hours[12]. Female mice were not used for these studies because of their documented ability to metabolize APAP without hepatic toxicity[13].

**Mouse organ collection.** Mice were anesthetized with isoflurane using a Sigma Delta Vaporizer (Penlon #52606). Upon deep induction, the thoracic cavity was opened to expose the beating heart, and 300-500 µL of blood was collected via cardiac puncture before perfusion. The perfusion solution (1X PBS) was administered for 10 min with a peristaltic pump (Bio-Rad Low-Pressure Chromatography Pump EP-1) at a flow rate of 2 mL/min. For the collection of organs from RiboTag mice, perfusion was conducted manually with a hand-operated syringe. The perfusion solution was supplemented with 100 ug/mL cycloheximide (CHX); 40 mL of CHX-supplemented perfusion solution was used per mouse for TRAP Samples.

**Serum collection.** 300-500 µL of blood was collected via cardiac puncture before perfusion. 150-250 µl of blood was placed in microtainer blood collection tubes containing a clot activator/SST™ Gel (Amber) (Tiger Medical: 365978). Blood was allowed to sit for 5 min and was centrifuged at 7,500 g for 5 min at 22°C. Serum was collected and stored at -80°C and used for subsequent ALT/AST Measurements.

**Hepatotoxicity assessment.** Mouse serum was subjected to alanine aminotransferase (ALT; IDEXX: 99-11040-01) and aspartate aminotransferase (AST; IDEXX: 99-11069-01) measurements using a Catalyst One Chemistry Analyzer from IDEXX Laboratories, Inc. (Westbrook, ME).

**Evans blue (EB) assay.** At 6 hr and 24 hr after APAP overdose, Evans blue dye (Sigma: E2129; 1% in 0.9% saline) was injected retro-orbitally into anesthetized mice at a dose of 4 µL/g body weight. The dye was allowed to circulate for 30 min without anesthesia. Mice were then re-anesthetized, and the left lateral lobes of the liver were harvested to assess dye leakage and permeability. Livers were then weighed and dried, and EB was extracted from the liver with formamide and normalized to the dry liver weight.

**Histological staining.** The left lateral lobes of mouse livers were fixed in 4% paraformaldehyde for 24 hr and embedded in paraffin blocks. Histological sections were cut at 5µm thickness using an HM 355S microtome from Microm International (Walldorf, Germany). Sections were mounted on positively charged UltraClear microscope slides from Denville Scientific Inc. (Holliston, MA) and allowed to dry overnight at 37°C. Hematoxylin and eosin (H&E) staining was performed as described previously [8], with a few modifications. Samples were washed for 5 min and were incubated in H&E for 45 seconds and 30 seconds, respectively.

**Microscopy and image acquisition.** Brightfield histological images were obtained with a Nikon DS-Fi1 camera coupled with an Eclipse 80i microscope from Nikon (Melville, NY) using 4x (NA 0.13) and 20x (NA 0.5) objectives.

**Endothelial Translating Ribosome Affinity Purification (TRAP).** We used a similar method for TRAP as was previously described [2], with some modifications. Frozen livers (25-100 mg) were pulverized using a BioPulverizer (Biospec Products: 59012N), with 750 µL of polysome buffer [50 mM Nuclease free Tris pH 7.5 (BioWorld: 21420063-3), 100 mM KCl (Sigma: P3911), 12 mM $MgCl_2$ (Sigma: M8266), 1% Igepal CA-630 (Sigma: 56741), 1 mM dithiothreitol (Sigma: 10197777001), 200 U/mL RNaseOUT (Thermo Fisher: 10777019), 1 mg/mL heparin sodium salt (Sigma: H3393), 100 µg/mL cycloheximide (Sigma: 01810) and an EDTA-free protease inhibitor cocktail (Sigma: 11836170001), buffer was prepared in DEPC-treated nuclease-free water (Thermo Fisher: AM9906)]. All liver samples were diluted in polysome buffers. Tissue homogenates were then centrifuged at 16,000 g at 4°C for 15 min, and the supernatant was transferred to a fresh tube and centrifuged again at 16,000 g at 4°C for 5 min. The protein concentration of the homogenate was measured using a detergent-compatible (DC) Protein Assay (Biorad: 5000111), and the homogenate was diluted to 5 mg/ml. 100 µl of homogenate was set aside to constitute the input fraction, against which the HA-enriched samples could be compared. To generate the EC-enriched samples, 400 µL of homogenate was then incubated with 5 µL (1:80) purified monoclonal rabbit anti-HA antibody (CST: C29F4) for 1 hour at 4°C under gentle rocking. 100 µL of magnetic protein G beads (New England BioLabs: S1430S) were then washed and equilibrated in 400 µL of the polysome buffer. The beads were then added to the antibody-incubated homogenates for an additional 30 min at 4°C under gentle rocking. The magnetic beads were subsequently washed 3 times with high salt buffer (polysome buffer with the KCl concentration adjusted to 300 mM). 500 µL of Trizol/chloroform was then added to the beads, and 100 µl of homogenate was set aside to constitute the input fraction. Total RNA from liver lysates and EC enriched fractions was isolated using a Qiagen RNeasy Mini Kit, including an on-column DNase digestion step, as described above, and quantified spectrophotometrically (NanoDrop). RNA was eluted with 30 uL of DEPC-treated nuclease-free water.

**Real-Time Quantitative PCR (qPCR).** Isolated RNA was converted to cDNA using the iScript cDNA Synthesis Kit (BioRad: 1708890). 100 ng of RNA was converted to cDNA in a 20 µL reaction volume. For TRAP-isolated samples, 10 µL of RNA was used in the reaction (~10-100 ng). Samples were incubated in the thermal cycler as per the manufacturer's instructions. Prepared cDNA was then diluted in DEPC-treated nuclease-free water with a final volume of 166 µL, i.e. 0.6 ng/µL of starting RNA. qPCR was then performed as previously described [8], using SsoAdvance Universal SYBR Green Supermix (BioRad: 1725275). qPCR was performed to validate TRAP samples before next-generation sequencing.

**mRNA library preparation.** RNA concentrations were measured using the Quant-it RiboGreen RNA kit (Thermo Fisher Scientific, catalog number R11490). RNA originating from either the total liver lysate (5 ng) or EC enriched fraction (10 ng) was converted into cDNA and subsequently amplified with the SMART-Seq v4 Ultra Low Input RNA kit (Takara Bio USA, catalog number 634891) according to the manufacturer's instructions. After bead purification (AMPure beads, Beckman Coulter, catalog number A63881), cDNA samples were sheared using the Covaris E220 Focused Ultra-sonicator, followed by Quant-iT PicoGreen quantification (Thermo Fisher Scientific, catalog number P11496). A total of 5 ng cDNA per sample was used for library preparation based on the KAPA Hyper Prep method (KAPA Biosystems, catalog number KK8504). Following NEXTflex adapter ligation (BIOO Scientific, catalog number 514104), libraries were size-selected based on a double bead purification prior to amplification. The concentration of post-amplified, bead-purified samples was measured using PicoGreen and the overall quality was assessed on the Agilent Tapestation 4200.

**Bioinformatics analysis.** 96 FASTQ Files containing reads on the TRAP data were generated by MedGenome. Reads were then aligned using HISAT2 and SALMON pipelines. HISAT2 generated reads to the genome and SALMON generated reads to the transcriptome. These reads were accumulated in a count matrix and a TPM matrix, respectively. The SALMON-generated TPM matrix files were used to identify TPM values for genes in the experimental samples, and different transcript variants were pooled to generate final TPM values using the aggregate function in R. The HISAT2-generated count matrix was used to generate 32 comparison files (using Deseq2 and EdgeR), consisting of relevant differentially expressed genes between our experimental groups. The Deseq2-generated results were used for the final analyses. Initial comparison files were generated after conducting alignment to 56,748 genes in the murine gene atlas to identify changes in relevant genes. All genes containing mean counts of less than 10 (averaged across all experimental groups) were removed. This resulted in comparison files containing 13,000-15,000 genes. The p values of these genes were rescaled using a Benjamini-Hochberg correction to account for the smaller gene set. To identify alterations in our endothelial-enriched samples, we then compared files from input vs EC fractions. This allowed for the identification of genes depleted in ECs (~3,000 genes), which are likely to be expressed in other liver cells. These genes were then removed from our datasets to allow sole focus on comparisons between ECs. This resulted in the EC datasets containing 10,000-12,000 genes. The p values of these genes were rescaled using a Benjamini-Hochberg correction to account for the smaller gene set.

**Structural modeling.** To identify the potential dimerization sites in the fourth transmembrane domains of murine PAR1 and PAR4, we generated *in silico* models of the receptors using SWISS-MODEL[14] and RaptorX[15], as described[16]. The murine model of PAR1 was generated by SWISS-MODEL, and the murine model of PAR4 was generated by RaptorX since SWISS-MODEL yielded a disjoined model of PAR4. Alignment of the receptor with existing models for PAR1 and PAR4, and correlation of likely dimerization residues with known dimerization residues in the human receptor[17], yielded a putative list of residues that likely mediate PAR1/4 dimerization in mice. For modeling the exodomain of PAR1, the exodomain that was generated by RaptorX was fused to the murine model of PAR1 generated by SWISS-MODEL and used to depict the *in silico* structure of murine PAR1 with an intact exodomain. PDB files for *in silico* models of both receptors will be provided upon request. Structural data were analyzed by UCSF Chimera and PyMOL.

**Statistical analysis.** All statistical analysis was performed in GraphPad Prism version 10.2.3 except for the bioinformatics data, which were performed in DESeq2 (as described in the "bioinformatics analysis" section). All bar and line graphs display results as mean $\pm$ SEM. All datasets include a minimum sample size of (n=3) biological replicates, which was sufficient for all inferential testing performed. Two-way ANOVA: For data that included two independent variables, a two-way ANOVA was used followed by a Tukey test to determine significant differences between groups. Given that we looked at changes in two independent variables, Tukey was chosen as a conservative test that balances type I and type II errors. Mixed effects model: This test was performed in place of a two-way ANOVA when analyzing unbalanced data (i.e., certain groups having more data points than others). Mixed models allow for unbalanced data to contribute to final results, whereas a two-way ANOVA cannot. When analyzing unbalanced data, a Benjamini, Krieger, and Yekutieli post-hoc test was chosen since it is less conservative than Tukey but still has significant power due to controlling the false discovery rate. For these analyses, q-values were displayed to show significance. Three-way ANOVA: This was performed for data that included three independent variables and was followed by a two-stage linear step-up procedure of Benjamini, Krieger, and Yekutieli for multiple comparisons testing to determine significant differences between groups. This post-hoc test was chosen since it is less conservative than Tukey but still has significant power due to controlling the false discovery rate. For these analyses, q-values were displayed to show significance. Welch t-test: This test was performed when analyzing data that included one independent variable, following verification that the data were normally distributed using a Shapiro-Wilk test. Multiple comparisons were performed by setting the desired false discovery rate (FDR) to 1% and performing a two-stage step-up method of Benjamini, Krieger, and Yekuteli. This post-hoc test was chosen since it is less conservative than Tukey but still has significant power due to controlling the false discovery rate.

Note that Shapiro-Wilk normality tests were performed on all data. Two- and three-way ANOVAs were still performed even when a single data point caused the normality test to fail since these ANOVAs can still generate valid outcomes as long as the dependent variables are approximately normally distributed for the compared groups according to the central limit theorem. This rationale was also applied to the mixed effects model as well; since this dataset included a large "n", the central limit theorem condones the use of non-normal data.

**Supplementary Appendix (online files)**

Supplemental File 1: (SF1.Pathways) List of altered IPA pathways in the liver and endothelium following APAP overdose.

Supplemental File 2: (SF2.TPMValues_Means) SALMON generated TPM values of the means of each group in our EC TRAPseq dataset (16 groups); 55,372 features annotated.

Supplemental File 3: (SF3.TPMValues) SALMON generated TPM values of all 48 samples generated by EC TRAPseq; 55,372 features annotated.

Supplemental File 4: (SF4.CountsFile) HISAT2 generated gene counts of all 48 samples generated by EC TRAPseq; 56,748 features annotated.

Supplemental File 5: (SF5.BiorenderLicenses) Copy of all biorender licenses for figures generated in the manuscript.

**Supplemental Table 1 (ST1): Genotyping Primers**

| Allele | Forward Primer | Reverse Primer | $T_m$ (C) | Bands (bp) |
|---|---|---|---|---|
| *Par1* | 5' - CATGGGGGAAGCTATCAGAA | 5' - CCAAAACCGAGTCCAAAAGA | 55 | WT – 202 Flox – 236 |
| *Par4* | 5' - GGGATGTTGTGGAGATTTGG | 5' - ATGGCTTCCCCTGGTACTCT | 55 | WT – 431 Flox – 465 |
| RiboTag | 5' - GGGAGGCTTGCTGGATATG | 5' - TTTCCAGACACAGGCTAAGTACAC | 55 | WT – 243 Flox – 290 |
| *Cdh5(PAC)-Cre$^{ERT2}$* | 5' - TCCTGATGGTGCCTATCCTC | 5' - CGAACCTGGTCGAAATCAGT | 55 | 473 |
| *Pf4-Cre* | 5' - CCCATACAGCACACCTTTTG | 5' -TGCACAGTCAGCAGGTT | 58 | 450 |

### Supplemental Table 2 (ST2): Description of statistical tests used for comparisons

| Figures | Panel | Normality | Equal Variance | Transformed | Statistical Test | Post hoc Test | Comparisons | q value | p value | n |
|---|---|---|---|---|---|---|---|---|---|---|
| 1 | C | NO (Shapiro-Wilk) | | | Mixed Effects Model (REML) | Benjamini, Krieger and Yekutieli | Control APAP 6 Hour v. *Par1/4*$^{iECko}$ APAP 6 Hour | 0.0203 | 0.0106 | n>8 |
| | | | | | | | Control APAP 24 Hour v. *Par1/4*$^{iECko}$ APAP 24 Hour | 0.0074 | 0.0141 | |
| | | | | | | | Control APAP 24 Hour v. *Par1*$^{iECko}$ APAP 24 Hour | 0.0335 | 0.0956 | |
| | | | | | | | Control APAP 24 Hour v. *Par4*$^{iECko}$ APAP 24 Hour | 0.0074 | 0.0072 | |
| 2 | A | NO (Shapiro-Wilk) | NO (Spearmans) | | Two-way ANOVA | Tukey | Control Saline v. Control APAP | | <0.0001 | n>6 |
| | | | | | | | *Par1/4*$^{iECko}$ Saline v. *Par1/4*$^{iECko}$ APAP | | 0.0095 | |
| | | | | | | | Control APAP v. *Par1/4*$^{iECko}$ APAP | | 0.0388 | |
| | B | NO (Shapiro-Wilk) | NO (Spearmans) | | Two-way ANOVA | Tukey | Control Saline v. Control APAP | | <0.0001 | n>4 |
| | | | | | | | *Par1*$^{iECko}$ Saline v. *Par1*$^{iECko}$ APAP | | 0.0015 | |
| | | | | | | | Control APAP v. *Par1*$^{iECko}$ APAP | | 0.0209 | |
| | C | NO (Shapiro-Wilk) | NO (Spearmans) | | Two-way ANOVA | Tukey | Control Saline v. Control APAP | | <0.0001 | n>4 |
| | | | | | | | *Par4*$^{iECko}$ Saline v. *Par4*$^{iECko}$ APAP | | 0.003 | |
| | | | | | | | Control APAP v. *Par4*$^{iECko}$ APAP | | 0.0426 | |
| | D | NO (Shapiro-Wilk) | NO (Spearmans) | | Two-way ANOVA | Tukey | Control Saline v. Control APAP | | <0.0001 | n>3 |
| | | | | | | | *Par4*$^{PLTko}$ Saline v. *Par4*$^{PLTko}$ APAP | | <0.0001 | |
| | | | | | | | Control APAP v. *Par4*$^{PLTko}$ APAP | | 0.9895 | |
| 3 | A | YES (Shapiro-Wilk) | NO (Spearmans) | | Two-way ANOVA | Tukey | Control Saline v. Control APAP | | <0.0001 | n>4 |
| | | | | | | | *Par1/4*$^{iECko}$ Saline v. *Par1/4*$^{iECko}$ APAP | | <0.0001 | |
| | | | | | | | Control APAP v. *Par1/4*$^{iECko}$ APAP | | 0.9308 | |
| | B | YES (Shapiro-Wilk) | NO (Spearmans) | | Two-way ANOVA | Tukey | Control Saline v. Control APAP | | <0.0001 | n>3 |
| | | | | | | | *Par1*$^{iECko}$ Saline v. *Par1*$^{iECko}$ APAP | | <0.0001 | |
| | | | | | | | Control APAP v. *Par1*$^{iECko}$ APAP | | 0.9166 | |
| | C | NO (Shapiro-Wilk) | NO (Spearmans) | | Two-way ANOVA | Tukey | Control Saline v. Control APAP | | 0.0017 | n>3 |
| | | | | | | | *Par4*$^{iECko}$ Saline v. *Par4*$^{iECko}$ APAP | | 0.0102 | |
| | | | | | | | Control APAP v. *Par4*$^{iECko}$ APAP | | 0.5888 | |
| | D | YES (Shapiro-Wilk) | NO (Spearmans) | | Two-way ANOVA | Tukey | Control Saline v. Control APAP | | <0.0001 | n=5 |
| | | | | | | | *Par4*$^{PLTko}$ Saline v. *Par4*$^{PLTko}$ APAP | | <0.0001 | |
| | | | | | | | Control APAP v. *Par4*$^{PLTko}$ APAP | | 0.9982 | |
| | E | NO (Shapiro-Wilk) | NO (Spearmans) | | Three-way ANOVA | Benjamini, Krieger and Yekutieli | Control APAP 6 Hour v. *Par1/4*$^{iECko}$ APAP 6 Hour | 0.6066 | 0.3209 | n>3 |
| | | | | | | | Control Saline 6 Hour v. Control APAP 6 Hour | 0.0061 | 0.0006 | |
| | | | | | | | Control APAP 24 Hour v. *Par1/4*$^{iECko}$ APAP 24 Hour | 0.7840 | 0.8301 | |
| | | | | | | | Control Saline 24 Hour v. Control APAP 24 Hour | 0.0125 | 0.0026 | |
| | F | NO (Shapiro-Wilk) | NO (Spearmans) | | Three-way ANOVA | Benjamini, Krieger and Yekutieli | Control APAP 6 Hour v. *Par1/4*$^{iECko}$ APAP 6 Hour | 0.6952 | 0.9932 | n>3 |
| | | | | | | | Control Saline 6 Hour v. Control APAP 6 Hour | 0.0015 | 0.0002 | |
| | | | | | | | Control APAP 24 Hour v. *Par1/4*$^{iECko}$ APAP 24 Hour | 0.6952 | 0.8756 | |
| | | | | | | | Control Saline 24 Hour v. Control APAP 24 Hour | 0.0332 | 0.0126 | |
| 5 | A | NO (Shapiro-Wilk) | NO (Spearmans) | | Two-way ANOVA | Tukey | *Par4: Par1*$^{ECwt}$ Saline Liver v. *Par1*$^{iEChet}$ Saline Liver | >0.9999 | | n=3 |
| | | | | | | | *Par4: Par1*$^{iEChet}$ Saline Liver v. *Par1*$^{iECko}$ Saline Liver | >0.9999 | | |
| | | | | | | | *Par4: Par1*$^{ECwt}$ Saline Liver v. *Par1*$^{iECko}$ Saline Liver | >0.9999 | | |
| | | | | | | | *Par4: Par1*$^{ECwt}$ Saline Liver v. *Par1*$^{ECwt}$ Saline EC | 0.0002 | | |
| | | | | | | | *Par4: Par1*$^{iEChet}$ Saline Liver v. *Par1*$^{iEChet}$ Saline EC | 0.0236 | | |
| | | | | | | | *Par4: Par1*$^{ECwt}$ Saline EC v. *Par1*$^{iEChet}$ Saline EC | 0.0795 | | |
| | | | | | | | *Par4: Par1*$^{iEChet}$ Saline EC v. *Par1*$^{iECko}$ Saline EC | 0.0156 | | |
| | | | | | | | *Par4: Par1*$^{ECwt}$ Saline EC v. *Par1*$^{iECko}$ Saline EC | 0.0001 | | |
| | B | YES (Shapiro-Wilk) | NO (Spearmans) | | Two-way ANOVA | Tukey | *Par4: Par4*$^{ECwt}$ Saline Liver v. *Par4*$^{iEChet}$ Saline Liver | 0.9967 | | n=3 |
| | | | | | | | *Par4: Par4*$^{iEChet}$ Saline Liver v. *Par4*$^{iECko}$ Saline Liver | >0.9999 | | |
| | | | | | | | *Par4: Par4*$^{ECwt}$ Saline Liver v. *Par4*$^{iECko}$ Saline Liver | 0.9967 | | |
| | | | | | | | *Par4: Par4*$^{ECwt}$ Saline Liver v. *Par4*$^{ECwt}$ Saline EC | 0.0062 | | |
| | | | | | | | *Par4: Par4*$^{iEChet}$ Saline Liver v. *Par4*$^{iEChet}$ Saline EC | 0.1135 | | |
| | | | | | | | *Par4: Par4*$^{ECwt}$ Saline EC v. *Par4*$^{iEChet}$ Saline EC | 0.3036 | | |
| | | | | | | | *Par4: Par4*$^{iEChet}$ Saline EC v. *Par4*$^{iECko}$ Saline EC | 0.2153 | | |
| | | | | | | | *Par4: Par4*$^{ECwt}$ Saline EC v. *Par4*$^{iECko}$ Saline EC | 0.0057 | | |
| | C | NO (Shapiro-Wilk) | NO (Spearmans) | | Three-way ANOVA | Benjamini, Krieger and Yekutieli | *Par4: Par1*$^{iEChet}$ Saline Liver v. *Par1*$^{iEChet}$ EC | 0.0015 | 0.0002 | n=3 |
| | | | | | | | *Par4: Par1*$^{iEChet}$ APAP Liver v. *Par1*$^{iEChet}$ APAP EC | 0.0078 | 0.0026 | |
| | | | | | | | *Par4: Par1*$^{iECko}$ Saline Liver v. *Par1*$^{iECko}$ EC | 0.0088 | 0.0042 | |
| | | | | | | | *Par4: Par1*$^{iECko}$ APAP Liver v. *Par1*$^{iECko}$ APAP EC | 0.1486 | 0.0885 | |
| | | | | | | | *Par4: Par1*$^{iEChet}$ Saline EC v. *Par1*$^{iEChet}$ APAP EC | 0.0078 | 0.0028 | |
| | | | | | | | *Par4: Par1*$^{iEChet}$ Saline EC v. *Par1*$^{iECko}$ Saline EC | 0.1642 | 0.1173 | |
| | | | | | | | *Par4: Par1*$^{iEChet}$ APAP EC v. *Par1*$^{iECko}$ APAP EC | 0.1771 | 0.1476 | |
| | | | | | | | *Par4: Par1*$^{iECko}$ Saline EC v. *Par1*$^{iECko}$ APAP EC | 0.6121 | 0.7287 | |
| | D | NO (Shapiro-Wilk) | NO (Spearmans) | | Three-way ANOVA | Benjamini, Krieger and Yekutieli | *Par1: Par4*$^{iEChet}$ Saline Liver v. *Par4*$^{iEChet}$ EC | 0.0001 | <0.0001 | n=3 |
| | | | | | | | *Par1: Par4*$^{iEChet}$ APAP Liver v. *Par4*$^{EChet}$ APAP EC | 0.0443 | 0.0264 | |
| | | | | | | | *Par1: Par4*$^{iECko}$ Saline Liver v. *Par4*$^{iECko}$ EC | 0.0002 | <0.0001 | |
| | | | | | | | *Par1: Par4*$^{iECko}$ APAP Liver v. *Par4*$^{iECko}$ APAP EC | 0.0027 | 0.001 | |
| | | | | | | | *Par1: Par4*$^{iEChet}$ Saline EC v. *Par4*$^{iEChet}$ APAP EC | 0.0041 | 0.0019 | |
| | | | | | | | *Par1: Par4*$^{iEChet}$ Saline EC v. *Par4*$^{iECko}$ Saline EC | 0.6664 | 0.7182 | |
| | | | | | | | *Par1: Par4*$^{EChet}$ APAP EC v. *Par4*$^{iECko}$ APAP EC | 0.1469 | 0.1082 | |
| | | | | | | | *Par1: Par4*$^{iECko}$ Saline EC v. *Par4*$^{iECko}$ APAP EC | 0.1469 | 0.1224 | |
| S3 | A | YES (Shapiro-Wilk) | YES (Spearmans) | | Two-way ANOVA | Tukey | Control v. *Par4*$^{iECko}$ | 0.0159 | | n>3 |
| | | | | | | | Control v. *Par1*$^{iECko}$ | 0.0112 | | |
| | | | | | | | Control v. *Par1/4*$^{iECko}$ | 0.0010 | | |
| | | | | | | | *Par1*$^{iECko}$ v. *Par4*$^{iECko}$ | 0.9866 | | |
| | | | | | | | *Par1*$^{iECko}$ v. *Par1/4*$^{iECko}$ | 0.7272 | | |
| | | | | | | | *Par4*$^{iECko}$ v. *Par1/4*$^{iECko}$ | 0.4514 | | |
| S3 | B | YES (Shapiro-Wilk) | YES (Spearmans) | | Two-way ANOVA | Tukey | *Par4*$^{iECko}$ v. *Par1/4*$^{iECko}$ | 0.1782 | | |
| S10 | D | YES (Shapiro-Wilk) | | | Welch T Test | Benjamini-Hochberg | *Par1*: Brain Parenchyma v. Brain EC | 0.1935 | 0.136 | n=3 |
| | | | | | | | *Par1*: Heart Parenchyma v. Heart EC | 0.0339 | 0.0009 | |
| | | | | | | | *Par1*: Kidney Parenchyma v. Kidney EC | 0.0267 | 0.0021 | |
| | | | | | | | *Par1*: Liver Parenchyma v. Liver EC | 0.0929 | 0.038 | |
| | | | | | | | *Par1*: Lung Parenchyma v. Lung EC | 0.0299 | 0.0042 | |
| | D | | | | | | *Par4*: Heart Parenchyma v. Heart EC | 0.031 | 0.0005 | |
| | | | | | | | *Par4*: Kidney Parenchyma v. Kidney EC | 0.0284 | 0.0028 | |
| S11 | | YES (Shapiro-Wilk) | | | Welch T Test | Benjamini, Krieger and Yekutieli | *PAR1*: Static vs Flow | 0.4007 | 0.7935 | n=3 |
| | | | | | | | *PAR4*: Static vs Flow | 0.0021 | 0.002 | |
| | | | | | | | *KLF2*: Static vs Flow | 0.0324 | 0.0482 | |
| | | | | | | | *KLF4*: Static vs Flow | 0.0001 | 0.0007 | |

**Supplemental Table 3 (ST4): List of mouse genders used for the generation of figure data**

| Figure | Condition | Males | Females |
|---|---|---|---|
| 1C | *Control* | 80 | |
| | *Par1*$^{iECko}$ | 15 | |
| | *Par4*$^{iECko}$ | 8 | |
| | *Par1/4*$^{iECko}$ | 17 | |
| 2A | Saline Control | 6 | |
| | Saline *Par1/4*$^{iECko}$ | 7 | |
| | APAP Control | 10 | |
| | APAP *Par1/4*$^{iECko}$ | 7 | |
| 2B | Saline Control | 5 | |
| | Saline *Par1*$^{iECko}$ | 4 | |
| | APAP Control | 7 | |
| | APAP *Par1*$^{iECko}$ | 4 | |
| 2C | Saline Control | 6 | |
| | Saline *Par4*$^{iECko}$ | 4 | |
| | APAP Control | 8 | |
| | APAP *Par4*$^{iECko}$ | 5 | |
| 2D | Saline Control | 8 | |
| | Saline *Par4*$^{PLTko}$ | 5 | |
| | APAP Control | 6 | |
| | APAP *Par4*$^{PLTko}$ | 3 | |
| 3A | Saline Control | 6 | |
| | Saline *Par1/4*$^{iECko}$ | 4 | |
| | APAP Control | 3 | |
| | APAP *Par1/4*$^{iECko}$ | 7 | |
| 3B | Saline Control | 4 | |
| | Saline *Par1*$^{iECko}$ | 4 | |
| | APAP Control | 3 | |
| | APAP *Par1*$^{iECko}$ | 7 | |
| 3C | Saline Control | 3 | |
| | Saline *Par4*$^{iECko}$ | 3 | |
| | APAP Control | 4 | |
| | APAP *Par4*$^{iECko}$ | 4 | |
| 3D | Saline Control | 5 | |
| | Saline *Par4*$^{PLTko}$ | 5 | |
| | APAP Control | 5 | |
| | APAP *Par4*$^{PLTko}$ | 5 | |
| 3E | Saline Control– 6 hr | 5 | |
| | Saline Control – 24 hr | 6 | |
| | APAP Control – 6 hr | 5 | |
| | APAP Control – 24 hr | 5 | |
| | Saline *Par1/4*$^{iECko}$ – 6 hr | 3 | |
| | Saline *Par1/4*$^{iECko}$ – 24 hr | 4 | |
| | Control *Par1/4*$^{iECko}$ – 6 hr | 3 | |
| | Control *Par1/4*$^{iECko}$ – 24 hr | 6 | |
| 3F | Saline Control– 6 hr | 5 | |
| | Saline Control – 24 hr | 5 | |
| | APAP Control – 6 hr | 5 | |
| | APAP Control – 24 hr | 5 | |
| | Saline *Par1/4*$^{iECko}$ – 6 hr | 3 | |
| | Saline *Par1/4*$^{iECko}$ – 24 hr | 4 | |
| | Control *Par1/4*$^{iECko}$ – 6 hr | 3 | |
| | Control *Par1/4*$^{iECko}$ – 24 hr | 6 | |
| 4B | Saline *Par4*$^{iEChet}$ Liver | 3 | |
| | Saline *Par4*$^{iECko}$ Liver | 3 | |
| | APAP *Par4*$^{iEChet}$ Liver | 3 | |
| | APAP *Par4*$^{iECko}$ Liver | 3 | |
| | Saline *Par1*$^{iEChet}$ Liver | 3 | |
| | Saline *Par1*$^{iECko}$ Liver | 3 | |
| | APAP *Par1*$^{iEChet}$ Liver | 3 | |
| | APAP *Par1*$^{iECko}$ Liver | 3 | |
| | Saline *Par4*$^{iEChet}$ EC | 3 | |
| | Saline *Par4*$^{iECko}$ EC | 3 | |
| | APAP *Par4*$^{iEChet}$ EC | 3 | |
| | APAP *Par4*$^{iECko}$ EC | 3 | |
| | Saline *Par1*$^{iEChet}$ EC | 3 | |
| | Saline *Par1*$^{iECko}$ EC | 3 | |
| | APAP *Par1*$^{iEChet}$ EC | 3 | |
| | APAP *Par1*$^{iECko}$ EC | 3 | |

| Figure | Condition | Males | Females |
|---|---|---|---|
| 4C | Saline *Par1*$^{iEChet}$ Liver | 3 | |
| | APAP *Par1*$^{iEChet}$ Liver | 3 | |
| 4D | Saline *Par1*$^{iEChet}$ EC | 3 | |
| | APAP *Par1*$^{iEChet}$ EC | 3 | |
| 5A | Saline *Par1*$^{ECwt}$ Liver | 3 | |
| | Saline *Par1*$^{iEChet}$ Liver | 3 | |
| | Saline *Par1*$^{iECko}$ Liver | 3 | |
| | Saline *Par1*$^{ECwt}$ EC | 3 | |
| | Saline *Par1*$^{iEChet}$ EC | 3 | |
| | Saline *Par1*$^{iECko}$ EC | 3 | |
| 5B | Saline *Par4*$^{ECwt}$ Liver | 3 | |
| | Saline *Par4*$^{iEChet}$ Liver | 3 | |
| | Saline *Par4*$^{iECko}$ Liver | 3 | |
| | Saline *Par4*$^{ECwt}$ EC | 3 | |
| | Saline *Par4*$^{iEChet}$ EC | 3 | |
| | Saline *Par4*$^{iECko}$ EC | 3 | |
| 5C | Saline *Par1*$^{iEChet}$ Liver | 3 | |
| | APAP *Par1*$^{iEChet}$ Liver | 3 | |
| | Saline *Par1*$^{iECko}$ Liver | 3 | |
| | APAP *Par1*$^{iECko}$ Liver | 3 | |
| | Saline *Par1*$^{iEChet}$ EC | 3 | |
| | APAP *Par1*$^{iEChet}$ EC | 3 | |
| | Saline *Par1*$^{iECko}$ EC | 3 | |
| | APAP *Par1*$^{iECko}$ EC | 3 | |
| 5D | Saline *Par4*$^{iEChet}$ Liver | 3 | |
| | APAP *Par4*$^{iEChet}$ Liver | 3 | |
| 6J | Saline *Par4*$^{iECko}$ Liver | 3 | |
| | APAP *Par4*$^{iECko}$ Liver | 3 | |
| 6I | Saline *Par4*$^{iEChet}$ EC | 3 | |
| | APAP *Par4*$^{iEChet}$ EC | 3 | |
| 6J | Saline *Par4*$^{iECko}$ EC | 3 | |
| | APAP *Par4*$^{iECko}$ EC | 3 | |
| 6A/C | APAP *Par1*$^{iEChet}$ EC | 3 | |
| | APAP *Par1*$^{iECko}$ EC | 3 | |
| 6B | APAP *Par4*$^{iEChet}$ EC | 3 | |
| | APAP *Par4*$^{iECko}$ EC | 3 | |
| 6D | Saline *Par1*$^{iEChet}$ EC | 3 | |
| | Saline *Par1*$^{iECko}$ EC | 3 | |
| | APAP *Par1*$^{iEChet}$ EC | 3 | |
| | APAP *Par1*$^{iECko}$ EC | 3 | |
| S3A | APAP Control | 27 | |
| | APAP *Par4*$^{iECko}$ | 5 | |
| | APAP *Par1*$^{iECko}$ | 4 | |
| | APAP *Par1/4*$^{iECko}$ | 6 | |
| S3B | APAP Control | 3 | |
| | APAP *Par4*$^{iECko}$ | 3 | |
| | APAP *Par1*$^{iECko}$ | 3 | |
| | APAP *Par1/4*$^{iECko}$ | 3 | |
| S5A | APAP *Par1*$^{iEChet}$ Liver | 3 | |
| | APAP *Par1*$^{iECko}$ Liver | 3 | |
| S5B | APAP *Par1*$^{iEChet}$ EC | 3 | |
| | APAP *Par1*$^{iECko}$ EC | 3 | |
| S6A/B | Saline *Par1*$^{iEChet}$ Liver | 3 | |
| | Saline *Par1*$^{iEChet}$ EC | 3 | |
| S7A-D | APAP *Par1*$^{iEChet}$ Liver | 3 | |
| | APAP *Par1*$^{iECko}$ Liver | 3 | |
| S8 | APAP *Par4*$^{iEChet}$ Liver | 3 | |
| | APAP *Par4*$^{iECko}$ Liver | 3 | |
| S9 | APAP *Par1*$^{iEChet}$ EC | 3 | |
| | APAP *Par1*$^{iECko}$ EC | 3 | |

**Supplemental Table 4 (ST4): Major Resources Table**

**Animals (in vivo studies)**

| Species | Vendor or Source | Background Strain | Sex | Persistent ID / URL |
|---|---|---|---|---|
| Mouse | Generated for our lab by ViewSolid Biotech, Inc | *Par1*$^{flox}$ | M | |
| Mouse | Generated for our lab by ViewSolid Biotech, Inc | *Par4*$^{flox}$ | M | |
| Mouse | Taconic | *Cdh5(PAC)-Cre*$^{ERT2}$ | M | #13073 |
| Mouse | Jackson Laboratory | *Pf4-Cre* | M | #008535 |
| Mouse | Jackson Laboratory | RiboTag | M | #029977 |

**Antibodies**

| Target antigen | Vendor or Source | Catalog # | Working concentration | Lot # | Persistent ID / URL |
|---|---|---|---|---|---|
| HA-Tag | CST | C29F4 | 1:80 (IP) | | RRID: AB_1549585 |

**Data & Code Availability**

| Description | Source / Repository | Persistent ID / URL |
|---|---|---|
| HISAT2 Pipeline | OMRF Center for Bioinformatics Data Science/Github | https://github.com/rrajala2/HISAT2_COMPARE.git |
| SALMON Pipeline | OMRF Center for Bioinformatics Data Science/Github | https://github.com/rrajala2/SALMON.git |
| R Code for Differential Expression | Rahul Rajala/Github | https://github.com/rrajala2/RCODE.git |
| FASTQ files | GSE266395/Gene Expresson Omnibus | to https://urldefense.com/v3/__https://www.ncbi.nlm.nih.gov/geo/query/acc.cgi?acc=GSE266395__;!!GNU8KkXDZlD12Q!7LVT74oN8sUWfg2ZC65PAyVrrpVYNL8mp2IiHPS1LII-XE5yGZ5CRdzd5vHb_77Y_a1weRb2Shlvr4gSd5YQ5B2C$ [ncbi[.]nlm[.]nih[.]gov] Enter token crelemwsflwxpur into the box |

**Other**

| Description | Source / Repository | URL / Catalog Number |
|---|---|---|
| **Software** | | |
| GraphPad Prism | GraphPad Software | https://www.graphpad.com/features |
| PyMOL | Schrodinger, LLC | https://pymol.org/ |
| UCSF Chimera | 2018 Regents of the University of California | https://www.cgl.ucsf.edu/chimera/ |
| RStudio Desktop | posit | https://posit.co/download/rstudio-desktop/ |
| R | 2024 The R Foundation for Statistical Computing | https://www.r-project.org/ |
| HISAT2 | UT Southwestern Lyda Hill Department of Bioinformatics | https://daehwankimlab.github.io/hisat2/ |
| SALMON | COMBINE-Labs | https://combine-lab.github.io/salmon/ |
| Adobe Photoshop 2024 | Adobe | https://www.adobe.com/products/photoshop/ |
| **Drugs/Agents/Kits/Equipment Used** | | |
| Tamoxifen | Sigma | T5648 |

| | | |
|---|---|---|
| Acetaminophen (APAP) | Sigma | A5000 |
| Cycloheximide (CHX) | Sigma | 01810 |
| Evans blue dye | Sigma | E2129 |
| Sigma Delta Vaporizer | Penlon | 52606 |
| Bio-Rad Low-Pressure Chromatography Pump EP-1 | BioRad | N/A |
| Serum Vial [clot activator/SST™ Gel (Amber)] | Tiger Medical | 365978 |
| ALT test clips | IDEXX | 99-11040-01 |
| AST test clips | IDEXX | 99-11069-01 |
| UltraClear microscope slides | Denville Scientific Inc | 14634019 |
| BioPulverizer | Biospec Products | 59012N |
| Nuclease free Tris pH 7.5 | BioWorld | 21420063-3 |
| KCl | Sigma | P3911 |
| $MgCl_2$ | Sigma | M8266 |
| Igepal CA-630 | Sigma | 56741 |
| dithiothreitol | Sigma | 10197777001 |
| RNaseOUT | Thermo Fisher | 10777019 |
| heparin sodium salt | Sigma | H3393 |
| EDTA-free protease inhibitor cocktail | Sigma | 11836170001 |
| DEPC-treated nuclease-free water | Thermo Fisher | AM9906 |
| Detergent-compatible (DC) Protein Assay | BioRad | 5000111 |
| RNeasy Mini Kit | Qiagen | 74106 |
| iScript cDNA Synthesis Kit | BioRad | 1708890 |
| SsoAdvance Universal SYBR Green Supermix | BioRad | 1725275 |
| Quant-it RiboGreen RNA kit | Thermo Fisher Scientific | R11490 |
| SMART-Seq v4 Ultra Low Input RNA kit | Takara Bio USA | 634891 |
| AMPure beads | Beckman Coulter | A63881 |
| E220 Focused Ultra-sonicator | Covaris | 500239 |
| Quant-iT PicoGreen quantification | Thermo Fisher Scientific | P11496 |
| KAPA Hyper Prep method | KAPA Biosystems | KK8504 |
| NEXTflex adapter ligation | BIOO Scientific, catalog number | 514104 |
| Agilent Tapestation 4200 | Agilent | G2991BA |
| Hematoxylin Stain Solution | Ricca chemical company | 3536-32 |
| Eosin Y | J.T.Baker | L088-03 |
